# The middle lipin (M-Lip) domain is a new dimeric protein fold that binds membranes

**DOI:** 10.1101/2021.03.19.436211

**Authors:** Weijing Gu, Shujuan Gao, Huan Wang, Kaelin D. Fleming, Reece M. Hoffmann, Jong Won Yang, Nimi M. Patel, Yong Mi Choi, John E. Burke, Karen Reue, Michael V. Airola

**Affiliations:** Department of Biochemistry and Cell Biology, Stony Brook University, Stony Brook NY 11794, USA; Department of Human Genetics, David Geffen School of Medicine at UCLA, Los Angeles CA 90095, USA; Department of Biochemistry and Microbiology, University of Victoria, Victoria BC V8N 1A1, Canada

## Abstract

Phospholipid synthesis and fat storage as triglycerides is regulated by lipin phosphatidic acid phosphatases (PAPs), whose enzymatic PAP function requires association with cellular membranes. Using hydrogen deuterium exchange mass spectrometry, we find mouse lipin 1 binds membranes through an N-terminal amphipathic helix and a middle lipin (M-Lip) domain that is conserved in mammalian and mammalian-like lipins. Crystal structures of the M-Lip domain reveal a previously unrecognized and novel protein fold that dimerizes. The isolated M-Lip domain binds membranes both in vitro and in cells through conserved basic and hydrophobic residues. Deletion of the M-Lip domain in full-length lipin 1 influences PAP activity, membrane binding, subcellular localization, oligomerization, and adipocyte differentiation, but does not affect transcriptional co-activation. This establishes the M-Lip domain as a new dimeric protein fold that binds membranes and is critical for full functionality of mammalian lipins.

## Introduction

Lipins are magnesium-dependent phosphatidic acid phosphatases (PAPs) that catalyze the dephosphorylation of the membrane lipid phosphatidic acid (PA) to produce diacylglycerol (DAG)^1^. The conversion of PA to DAG by lipins regulates de novo phospholipid biosynthesis^2^, cellular signaling^3^, chylomicron biogenesis^4^, adipocyte differentiation^5^, and fat storage as triglycerides^6^. Mutations that reduce lipin PAP activity^7, 8, 9^ are associated with metabolic diseases including rhabdomyolysis^10^, Majeed syndrome^11^, lipodystrophy^6^, statin-induced myopathy^12, 13^, and insulin resistance^6, 14^. Mammalian lipins also function as transcriptional co-activators to affect the transcription of genes for fatty acid oxidation^8, 15^. There are three mammalian lipin paralogs: lipin 1, lipin 2 and lipin 3^6, 16^. Many insights into lipin PAP function have come from studies of the yeast lipin homolog, *S. cerevisiae* PA phosphohydrolase 1 (*Sc* Pah1)^1, 17^.

The architecture of mammalian lipins and *Sc* Pah1 differ, but all lipin/Pah homologs share two common and conserved regions called the N-Lip and C-Lip regions, which are located at the respective N- and C-termini of mammalian lipins^6^. Recently, we determined the structure of *Tetrahymena thermophila* Pah2 (*Tt* Pah2), which revealed the N-Lip and C-Lip regions co-fold to form a catalytic unit comprised of a split immunoglobulin-like (Ig-like) domain and a haloacid dehalogenase-like (HAD-like) catalytic domain^18^. The N-Lip and C-Lip regions are separated by an extended linker that varies in length and sequence across species^17, 19^. In mammalian lipins, this linker is 500 amino acids and can be hyperphosphorylated^7, 20, 21, 22^, sumoylated^23^, or acetylated^24^. Phosphorylation, sumoylation, or acetylation within the linker region is reported to regulate the subcellular location and activity of mammalian lipins^7, 21, 22, 23, 24, 25^. However, it is not known if the linker region has additional roles.

As the only enzyme in the glycerol-3-phosphate pathway that is not constitutively membrane-bound, the regulation of lipin/Pah membrane association is a determinant of its enzyme activity. In vitro, purified *Sc* Pah1, *Tt* Pah2, and mammalian lipins are recruited to membranes containing their substrate, PA^18, 22, 26^. Lipin/Pahs lack canonical lipid binding domains (e.g., PH and PX domains) found in other membrane-binding proteins, but contain a conserved N-terminal amphipathic helix that is necessary for lipin/Pahs to bind membranes in vitro and in cells^18, 26^ In addition, a nuclear localization signal/polybasic region in mammalian lipins has been implicated in membrane binding^27, 28^.

We sought to characterize how lipins bind membranes and in the process identified a new middle lipin (M-Lip) domain that is universally conserved in mammalian and mammalian-like lipins, but not present in *Sc* Pah1. Herein, we report the structural and functional characterization of the M-Lip domain. Our principal findings are that the M-Lip domain is a new protein fold that forms a dimer, binds membranes, and can affect lipin PAP activity, oligomerization, subcellular localization, and adipogenesis.

## Results

### Structure and dynamics of lipin 1

Full-length mouse lipin 1α (herein referred to as lipin 1) was purified from Sf9 cells and the structure and dynamics were probed using hydrogen deuterium exchange mass spectrometry (HDX-MS). HDX-MS measures the exchange rate of amide hydrogens with deuterium. The major determinant of exchange is the stability of secondary structure^29^. Thus, HDX-MS provides a readout of secondary structure dynamics. In addition, HDX experiments with extremely short exposures of D_2_O can be used to identify disordered regions within proteins when compared to a fully deuterated condition^29^.

HDX experiments were carried out with a short pulse of deuterium exposure to map regions of order/disorder in full-length lipin 1. A total of 189 peptides spanning 83.6% of the primary sequence were identified and quantified. Peptides for residues 121-256 and residue 413 were not identified by tandem MS/MS. Residues with no MS coverage were all located between the N-Lip and C-Lip regions, and near the highly basic nuclear localization signal.

The N-Lip and C-Lip regions both had low rates of deuterium exchange with a 3 sec pulse of D_2_O exposure at 4°C, which indicates these regions were largely ordered into secondary structure elements **(Fig. 1b)**. This suggests that the N-Lip and C-Lip regions co-fold to form the split Ig-like domain and HAD-like domain observed in the *Tt* Pah2 structure^18^, and that in vitro their interaction is most likely constitutive.

**Figure 1.**
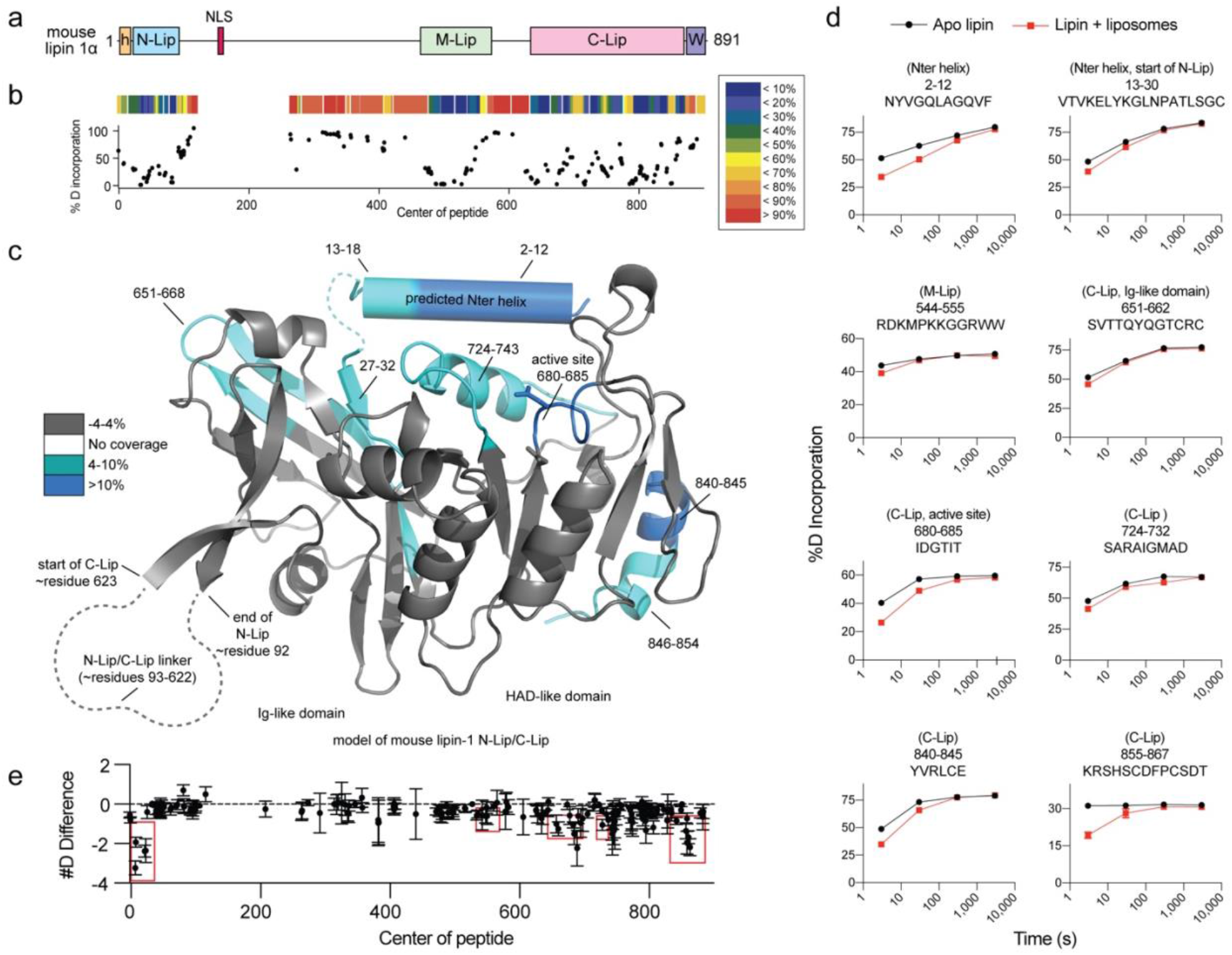
Structure and dynamics of lipin 1. **a**. Domain architecture of mouse lipin 1 drawn to scale. Mammalian lipins conserve three regions/domains: the N-Lip, M-Lip, and C-Lip. The N-Lip and C-Lip are predicted to co-fold to form two domains: an Ig-like and HAD-like domain. The positions of the N-ter amphipathic helix (orange h), nuclear localization signal (NLS), and conserved Trp-motif (purple W) are indicated. **b**. % deuterium incorporation after a 3 sec deuterium exposure in the absence of liposomes. Each point represents a single peptide, with them being graphed on the x-axis according to their central residue. A heat map above is color coded according the legend. **c**. Regions of lipin 1 that showed significant decreases in exchange (defined as >4%, >0.4 Da, and a student t-test p<0.01) in the presence of liposomes are colored in blue according to the legend and shown on a model of the N-Lip and C-Lip regions generated by Phyre2. **d**. % deuterium incorporation of selected peptides at various time points (3, 30, 300, and 3000 seconds) in the absence and presence of liposomes. The error bars represent standard deviation (n=3), most are smaller than the size of the point. **e**. The sum of the # of deuterons protected from exchange in the presence of liposomes across all timepoints is shown. Each point represents a single peptide, with them being graphed on the x-axis according to their central residue. Error bars represent standard deviation (n=3).

The majority of the >500 residues that separate the N-Lip and C-Lip regions had high rates of deuterium exchange, indicating that they are likely disordered **(Fig. 1b)**. One exception was a continuous stretch of ∼100 residues that were protected from exchange, which is indicative of secondary structure formation **(Fig. 1b)**. This ordered region is called the middle lipin (M-Lip) domain and we discuss M-Lip in more detail below.

### Lipin 1 association with membranes

In line with previous observations^22, 27^, recombinant lipin 1 bound strongly to PC/PA liposomes (see below). To identify the membrane-binding regions of lipin 1 we employed HDX-MS, which has been particularly useful in examining protein-lipid interactions^30, 31^. HDX-MS experiments were carried out in the presence and absence of PC/PA liposomes and at 4 different time points of exchange (3, 30, 300, and 3000 sec).

HDX-MS revealed multiple peptides that were protected from H/D exchange in the presence of liposomes, which suggests these regions associate with membranes. The regions protected by membrane were distributed throughout the primary structure and clustered into several key areas **(Fig. 1c, d, e)**. This included (i) the N-terminus (residues 2-12, 13-30) that is predicted to form an amphipathic helix, (ii) the C-terminal end of the M-Lip domain (residues 544-555) that is enriched in basic and hydrophobic residues **(Fig. 2a)**, (iii) peptides in the Ig-like domain (residues 651-662) that are predicted to lie at the membrane interface, (iv) the catalytic active site of the HAD-like domain (residues 660-685, 724-732) where PA hydrolysis occurs, and (v) the C-terminal end of the C-Lip (residues 840-845, 855-867) that is situated between the end of the HAD-like domain and the conserved Trp motif^32^ **(Fig. 1a, c, d, e)**. As stated above, peptides containing the NLS were not identified by tandem MS/MS. Thus, we were unable to assess the dynamics of membrane association for the NLS, which has previously been implicated in lipin 1 membrane binding^27^.

**Figure 2.**
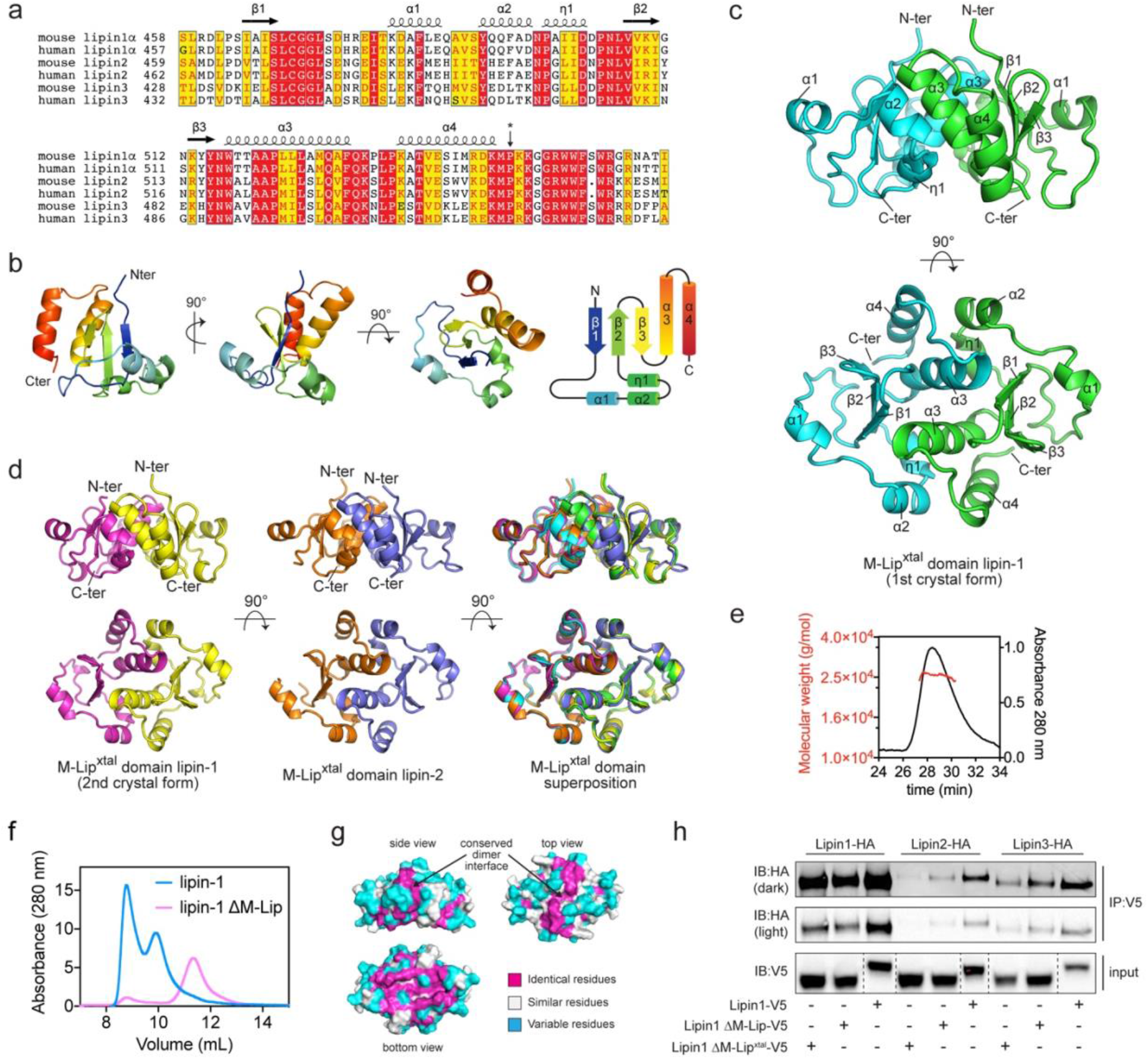
The M-Lip domain is a new protein fold that dimerizes. **a**. Sequence alignment of human and mouse M-Lip domains with secondary structure elements shown above. The asterisk indicates the last residue of the M-Lip^xtal^ domain. Red and yellow boxes indicate identical and positive homology residues. **b**. Structure of a single subunit of the lipin 1 M-Lip^xtal^ domain in three different views. The N- and C-termini are colored blue and red, respectively. **c**. The M-Lip^xtal^ domain forms a dimer with the α3 helices mediating the majority of contacts at the dimer interface. Individual subunits are colored green and cyan. **d**. Crystal structures and superimposition of the near identical lipin 1 and lipin 2 M-Lip^xtal^ domains. **e**. MALS data (left axis) with SEC traces (right axis) for the lipin 1 M-Lip^xtal^ domain reports a MW of 25 kDa, consistent with a dimer (MW of 26 kDa). **f**. SEC profiles of full-length and lipin 1 ΔM-Lip on SEC. Deletion of the M-Lip shifts the elution profile to a smaller apparent MW. **g**. Surface views of the lipin 1 M-Lip^xtal^ domain showing conservation of the dimer interface. **h**. Co-immunoprecipitation of HA-tagged lipins with V5-tagged lipin 1 constructs. Hepa1-6 cells were co-transfected with HA-tagged lipin 1, -2, and −3 and either V5-tagged lipin 1, ΔM-Lip, or ΔM-Lip^xtal^. Deletion of the ΔM-Lip or ΔM-Lip^xtal^ domains reduced the amount of HA-tagged lipin proteins that co-immunoprecipitated.

Notably, the membrane protected regions within the N-Lip and C-Lip regions of lipin 1 **(Fig. 1c)** were nearly identical with those previously observed for *Tt* Pah2^18^. This suggests the catalytic core of PAP enzymes utilize a conserved mechanism for membrane binding that involves an N-terminal amphipathic helix, the HAD-like active site, and portions of the Ig-like domain.

### The M-Lip domain

We next turned our attention to M-Lip, as the HDX-MS experiments suggested it may represent a third domain in lipin 1 that is involved in membrane binding. BLAST searches revealed that the M-Lip domain was selectively found in lipin homologs and was absolutely conserved in all mammalian lipins **(Fig. 2a)**. The M-Lip domain was also detected in some plant (*A. thaliana*), fungal (*C. neoformans*), ciliate (*T. thermophila*), insect (*D. melanogaster*), and apicomplexan (*P. falciparum*) lipin homologs **(Supplementary Fig. 1)** but was not detected in *Sc* Pah1. The M-Lip domain is thus one feature that distinguishes mammalian and mammalian-like lipin PAPs from *Sc* Pah1.

### Structure of the M-Lip domain reveals a new protein fold

The M-Lip domain did not share sequence homology with any domains of known function. To determine if M-Lip was indeed a protein domain with a defined tertiary structure, we sought to determine its structure. The mouse lipin 1 M-Lip domain was purified from *Escherichia coli*. The detergent Triton X-100 was required to prevent the M-Lip domain from pelleting during centrifugation after cell lysis but was not required in subsequent purification steps if high salt concentrations were maintained. Despite extensive efforts, we have yet to successfully crystallize the complete M-Lip domain.

We therefore truncated the M-Lip domain of mouse lipin 1 to remove the C-terminal cluster of hydrophobic and basic residues **(Fig. 2a)** implicated in membrane binding by HDX-MS **(Fig. 1d)**. We refer to this construct as the M-Lip^xtal^ domain. The M-Lip^xtal^ domain could be purified without detergent and was extremely stable with a melting temperature of 65 °C **(Supplementary Fig. 2)**. The structure of M-Lip^xtal^ domain of mouse lipin 1 was determined to resolutions of 1.5Å and 2.1Å in two unique space groups. Phases were obtained using single-wavelength anomalous diffraction from selenomethionine (SeMet)-derivatized protein **(Table 1)**. We also determined the structure of the mouse lipin 2 M-Lip^xtal^ domain to 2.4 Å resolution. The M-Lip^xtal^ domain from mouse lipin 3 was easily purified but did not crystallize.

**Table 1.**
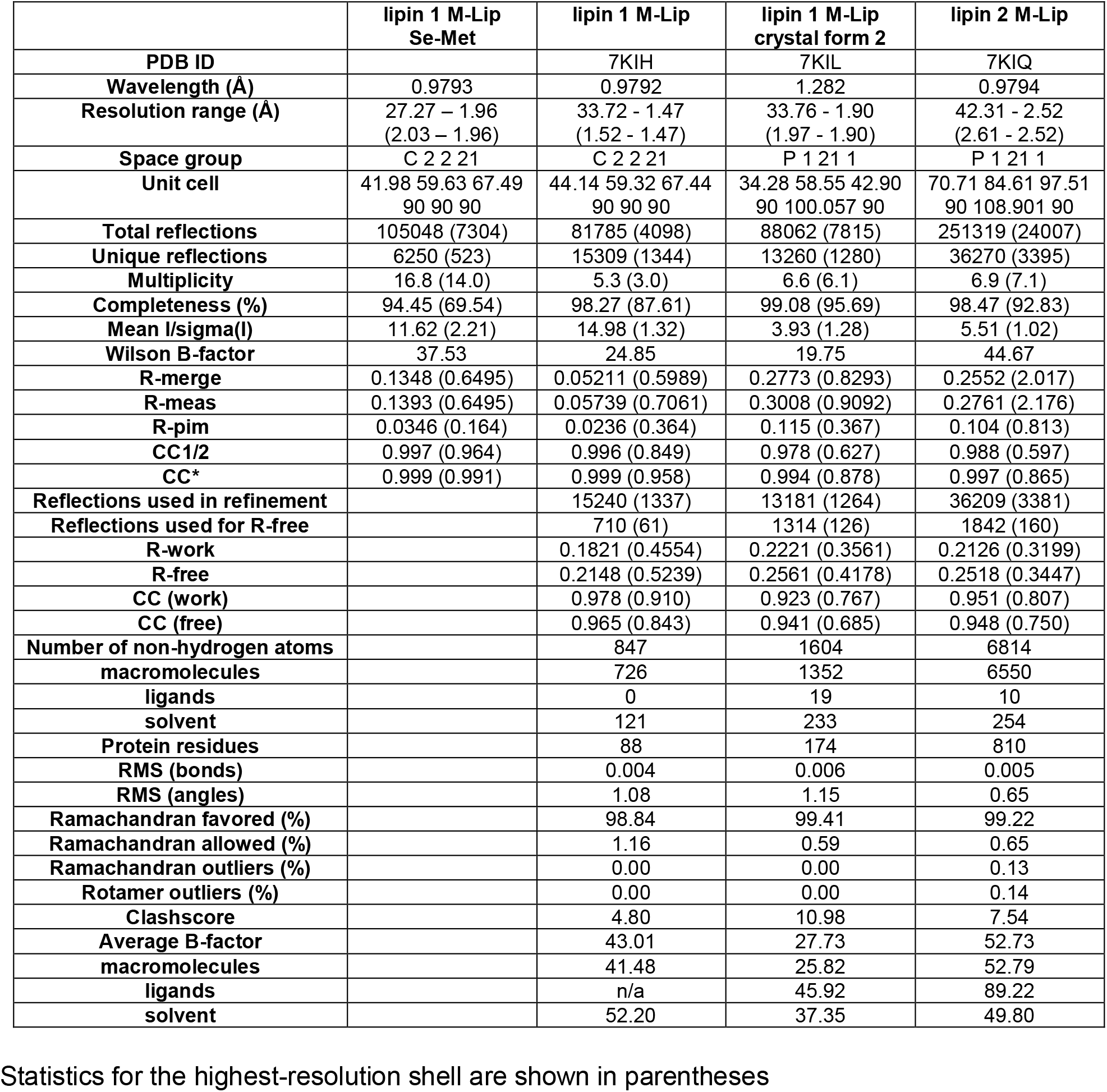
Data collection and refinement statistics.

The structure of the M-Lip^xtal^ domain revealed a protein fold with a three-stranded anti-parallel β-sheet at the core. Flanking the β-sheet were a set of two alpha helices (α1 and α2) and a short 3-10 helix (η1) that were oriented perpendicular to the β-sheet and two C-terminal α-helices (α3 and α4) that were oriented parallel to the β-sheet **(Fig. 2b)**. A Dali search^33^ did not identify any similar existing protein structures. Thus, the M-Lip domain represents a previously unrecognized and novel protein fold.

### The M-Lip domain forms a dimer

The M-Lip domain of mouse lipin 1 and lipin 2 formed a symmetric or near symmetric dimer in all three crystal forms with a root mean square deviation for all atoms ranging between 0.36 – 0.77 Å **(Fig. 2c, 2d)**. The observation of near identical dimers in three different crystal structures strongly suggested the native quaternary structure of the M-Lip domain as dimeric. To confirm this, we used size-exclusion chromatography coupled to multi-angle light scattering (SEC-MALS) to calculate the molecular weight (MW) of the M-Lip^xtal^ domain in solution. SEC-MALS reported a MW of 25 kDa for the mouse lipin 1 M-Lip^xtal^ domain, which was consistent with the MW of 26 kDa for a M-Lip^xtal^ dimer **(Fig. 2e)**.

Mammalian lipins are known to form both homo and hetero-oligomers^34^. We hypothesized that the M-Lip domain may be involved in lipin oligomerization. We therefore deleted the M-Lip domain in the context of full-length mouse lipin 1 (ΔM-Lip lipin 1) and purified ΔM-Lip lipin 1 from Sf9 cells. In comparison to wild-type lipin 1, the size exclusion profile of ΔM-Lip lipin 1 was shifted towards a lower molecular weight **(Fig. 2f)**. This is consistent with a role for the M-Lip domain in lipin 1 dimerization.

Analysis of the dimer interface of the M-Lip revealed an extensive network of interactions between the two subunits with residues within the α3 helix, which was located in the center of the dimer, mediating the majority of these interactions **(Fig. 2c)**. Notably, the residues involved in dimerization were highly conserved among human and mouse lipin 1, lipin 2, and lipin 3 paralogs **(Fig. 2a, h)**. Thus, we suspected that the M-Lip domain might also be involved in lipin hetero-oligomerization. Consistent with this hypothesis, deletion of the complete M-Lip domain or the M-Lip^xtal^ domain reduced the ability of lipin 1 to co-immunoprecipitate with lipin 1, lipin 2 and lipin 3 **(Fig. 2i)**.

### The M-Lip is not necessary for lipin transcriptional co-activator activity

The high conservation of the M-Lip region among mammalian lipins suggested that it plays a role(s) in lipin function. Mammalian lipins function both as PAP enzymes and as transcriptional co-activators for PPARα and PGC-1α with an LxxLL motif in the C-Lip critical for this latter function^15^. Since *Sc* Pah1 has neither an M-Lip domain nor transcription co-activator activity, we hypothesized M-Lip may be necessary for lipin co-activation function.

Using HEK293 cells and the established luciferase-based assay^8, 15^, lipin 1 increased the transcription of luciferase under the control of three peroxisome proliferator response elements (PPREs) in the presence of peroxisome-proliferator-activated receptor alpha (PPARα) and retinoic X receptor alpha (RXRα) **(Fig. 3a)**. Transcription was further increased in the presence of peroxisome proliferator-activated receptor gamma co-activator 1-alpha (PGC1α) **(Fig. 3a)**. Deletion of either the M-Lip or M-Lip^xtal^ sequences had no significant effect on PGC1α co-activation **(Fig. 3a)**. We concluded that the M-Lip domain is not necessary for lipin 1 to function as a transcriptional co-activator.

**Figure 3.**
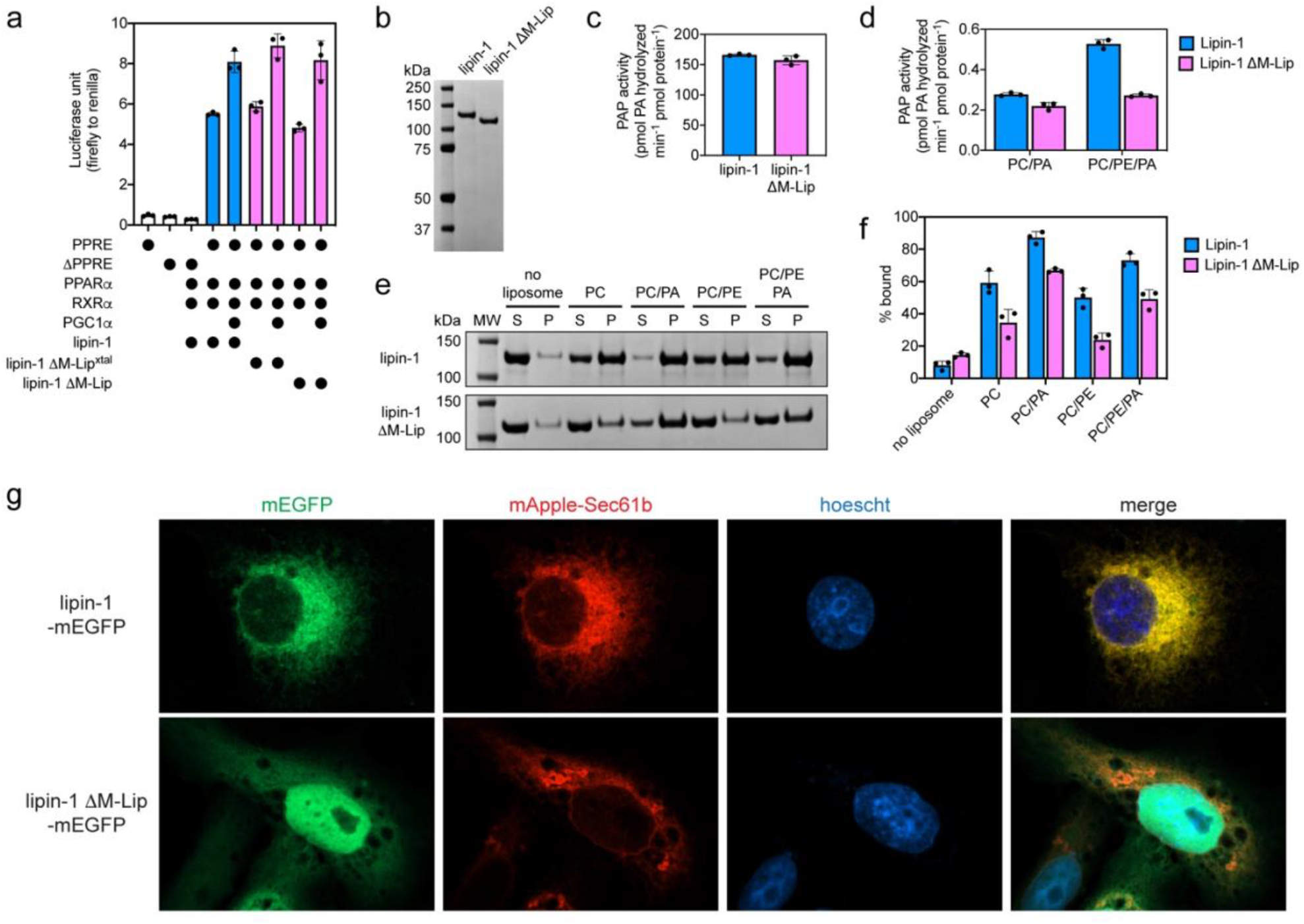
The M-Lip domain influences lipin 1 membrane binding and PAP activity. **a**. Deletion of the M-Lip or M-Lip^xtal^ domain does not affect lipin 1 transcriptional co-activation. HEK293 cells were co-transfected with lipin 1 or lipin 1 M-Lip domain deletion constructs with the transcription factors PPARα and RXRα, and the transcriptional co-activator PGC1-α. Transcriptional co-activation was quantitated by measuring firefly luciferase activity, which is under the control of three peroxisome-proliferation response elements (PPREs). Firefly luciferase activity was normalized to Renilla luciferase from a control plasmid that was included in each transfection. Data are the means and SDs of three experiments (n=3). **b**. SDS-PAGE of purified lipin 1 and lipin 1 ΔM-Lip used in PAP activity and liposome sedimentation assays. **c**. PAP activity of lipin 1 and lipin 1 ΔM-Lip in Triton X-100 mixed micelles with 10 mol% NBD-PA. Data are the means and SDs of three experiments (n=3). **d**. PAP activity of lipin 1 and lipin 1 ΔM-Lip in 90 mol% PC or 70/20 mol% PC/PE liposomes with 10 mol% NBD-PA. PC, phosphatidylcholine; PE, phosphatidylethalonine; PA, phosphatidic acid. Data are the means and SDs of three experiments (n=3). **e**. SDS-PAGE analysis of a liposome sedimentation assay reveals lipin 1 preferentially associates with liposomes containing 20 mol% PA. Deletion of the M-Lip domain (lipin 1 ΔM-Lip) results in a reduction of membrane association. S, supernatant; P, pellet; MW, molecular weight markers. **f**. Quantification of liposome association for lipin 1 and lipin 1 ΔM-Lip. Data are the means and SDs of three experiments (n=3). **g**. Confocal microscopy images of Cos-7 cells transiently transfected with monomeric enhanced GFP (mEGFP) fusions of either lipin 1 or lipin 1 ΔM-Lip (green) and the ER marker mApple-Sec61b (red). Hoechst stain (blue), nucleus.

### Deletion of the M-Lip domain influences lipin PAP activity

We next tested whether deletion of the M-Lip domain affected lipin 1 PAP activity in vitro using purified protein and the fluorescent substrate nitrobenzoxadiazole-phosphatidic acid (NBD-PA). Wild-type and ΔM-Lip lipin 1 had near identical PAP activities when the substrate NBD-PA was incorporated into Triton X-100 mixed micelles **(Fig. 3c)**.

We next assessed PAP activity with NBD-PA incorporated into liposomes, which represent a more physiologically relevant in vitro system. Liposomes composed solely of phosphatidylcholine (PC) or a mixture of PC and phosphatidylethanolamine (PE) were used, as the membrane lipid PE has previously been shown to increase PAP activity of lipin 1^22, 28^. The PAP activity of wild-type and ΔM-Lip lipin 1 were similar in PC liposomes **(Fig. 3d)**. However, unlike wild-type lipin 1, the activity of ΔM-Lip lipin 1 did not increase with the addition of PE in PC/PE liposomes **(Fig. 3d)**. Thus, PAP activity in PC/PE liposomes was ∼50% lower for ΔM-Lip lipin 1 compared to wild-type, which suggests that the M-Lip domain is involved in the PE-mediated effects on lipin 1 PAP activity.

### Deletion of M-Lip affects membrane binding and cellular localization

Given our findings that M-Lip influences PAP activity on PC/PE liposomes, we hypothesized that the M-Lip domain influences lipin 1 binding to membranes. Using a liposome sedimentation assay, we found that lipin 1 bound to liposomes containing the neutral lipids PC or PC/PE, and liposome association was further enhanced by the presence of 20 mol% PA **(Fig. 3e, f)**. Deletion of the M-Lip domain resulted in a consistent reduction of liposome association regardless of the liposome lipid composition **(Fig. 3e, f)**.

To assess if the M-Lip domain affects the ability of lipin 1 to localize to membranes in cells, we transiently transfected Cos-7 cells with wild-type and ΔM-Lip lipin 1 fused at their C-terminus with monomeric enhanced GFP (mEGFP) and assessed their subcellular localization using confocal microscopy. Lipin 1 displayed strong co-localization with the ER marker mApple-Sec61b. In contrast, ΔM-Lip lipin 1 accumulated in the nucleus and did not clearly co-localize with any membrane structures. We concluded that the M-Lip domain is necessary for proper subcellular localization and full-membrane binding in vitro.

### The isolated M-Lip domain binds membranes

To verify that the M-Lip domain directly binds membranes, we conducted liposome sedimentation assays using the isolated M-Lip and M-Lip^xtal^ domains purified from *E. coli*. The M-Lip domain bound to PC liposomes, and membrane association was greatly enhanced by the presence of the anionic lipids PA, phosphatidylserine, and phosphatidylinositol **(Fig. 4a, b)**. In contrast, the M-Lip^xtal^ domain, which lacks the conserved C-terminal hydrophobic and basic residues, exhibited weak association with liposomes that was not significantly influenced by the presence of anionic lipids **(Fig. 4a, b)**.

**Figure 4.**
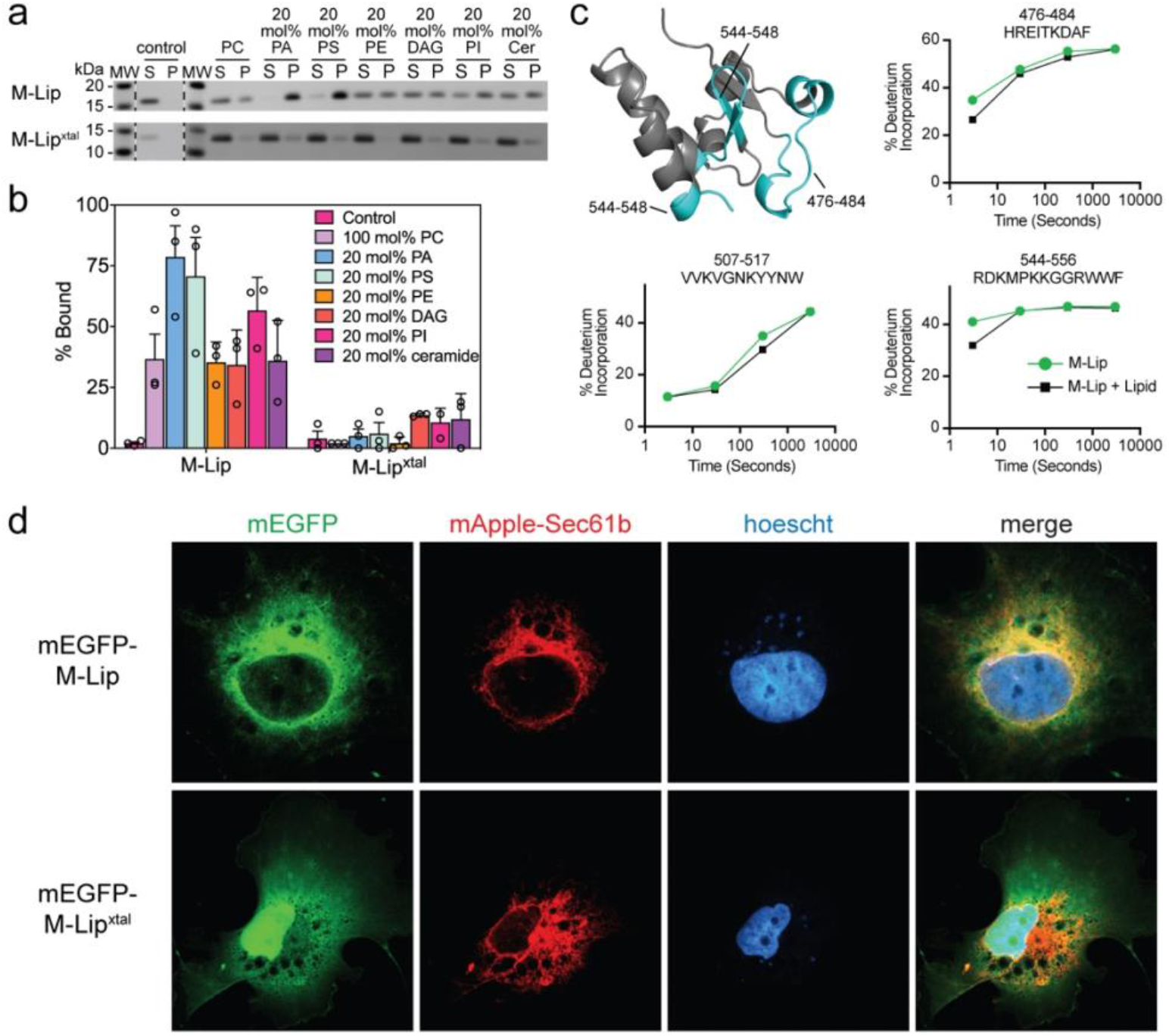
The M-Lip domain binds membranes in vitro and in cells. **a**. SDS-PAGE analysis of a liposome sedimentation assay reveals the M-Lip domain binds membranes and preferentially associates with liposomes containing anionic lipids (PA, PS, and PI). The M-Lip^xtal^ domain binds weakly to membranes. S, supernatant; P, pellet; MW, molecular weight markers; PC, phosphatidylcholine; PE, phosphatidylethalonine; PA, phosphatidic acid; PS, phosphatidylserine; DAG, diacylglycerol; PI, phosphatidylinositol; Cer, ceramide. **b**. Quantification of liposome association for the M-Lip and M-Lip^xtal^ domains. Data are the means and SDs of three experiments (n=3). **c**. Regions that showed significant decreases in exchange (defined as >5%, >0.4 Da, and a student t-test p<0.01) in the presence of liposomes are colored in blue according to the legend and displayed on a single subunit of the lipin 1 M-Lip^xtal^ domain. % deuterium incorporation of selected peptides at various time points (3, 30, 300, and 3000 seconds) in the absence and presence of liposomes. The error bars represent standard deviation (n=3), most are smaller than the size of the point. **d**. Confocal microscopy images of Cos-7 cells transiently transfected with monomeric enhanced GFP (mEGFP) fusions of either the M-Lip or M-Lip^xtal^ domains (green) and the ER marker mApple-Sec61b (red). Hoechst stain (blue), nucleus.

Next, we employed HDX-MS to identify the regions of the M-Lip domain that interact with membranes in an unbiased manner. HDX-MS experiments were conducted in the presence and absence of PC/PA liposomes. Several peptides were protected from deuterium exchange in the presence of liposomes. The strongest protection was observed for peptides containing a WWF motif, which are part of the conserved hydrophobic and basic residues at the C-terminal end of the M-Lip **(Fig. 2a, 4c)**. Two additional peptides present in the crystallized M-Lip^xtal^ domain were also protected **(Fig. 4c)**. These peptides mapped to the same surface, which suggests that the core of the M-Lip^xtal^ domain can also interact with membranes but is not the major determinant of membrane binding **(Fig. 4c)**.

To determine if the isolated M-Lip domain localized to membranes within cells, Cos-7 cells were transfected with M-Lip fusions with mEGFP. A mEGFP M-Lip fusion co-localized with the ER marker mApple-Sec61b **(Fig. 4d, Supplementary Fig. 4a)**. In contrast, the mEGFP M-Lip^xtal^ fusion accumulated in the nucleus with a minor co-localization with the ER, suggesting a role for the hydrophobic and basic residues at the C-terminus of M-Lip in membrane targeting **(Fig. 4d, Supplementary Fig. 4b)**. Taken together, these results identify the M-Lip domain as a new type of membrane binding domain.

### M-Lip role in adipogenesis

Lastly, we sought to characterize the role of the M-Lip domain in lipin 1 function in adipocytes. *Lpin1* is expressed in a bi-phasic manner during adipocyte differentiation, and has specific roles in both the early stages of adipogenesis and in the formation of mature, lipid-laden adipocytes^5^. In preadipocytes, lipin 1 PAP activity regulates PA signaling to promote expression of a key adipogenic transcription factor, peroxisome proliferator-activated receptor γ (PPARγ)^5, 35^. Lipin 1 PAP activity is also required in mature adipocytes for triglyceride synthesis and lipid hydrolysis^35, 36^. We investigated whether the M-Lip domain is required for the optimal activity of lipin 1 in the regulation of gene expression and lipid accumulation during adipocyte differentiation.

3T3-L1 preadipocytes were transfected with expression vectors for wild-type or ΔM-Lip lipin 1. We titrated expression levels of wild-type and ΔM-Lip lipin 1 to ensure that they were expressed at comparable levels **(Supplemental Fig. 5a)**. Expression of adipocyte genes was monitored at intervals during differentiation, and neutral lipid accumulation was examined after 5 days **(Fig. 5a)**. Wild-type lipin 1 expression, but not ΔM-Lip 1 expression, increased the levels of oil red O-stained lipids beyond those observed in cells transfected with a vector control when assessed at day 5 **(Fig. 5b and Supplemental Fig. 5b)**. Both wild-type and ΔM-Lip lipin 1 expression led to enhanced *Pparg* expression compared to vector controls, but ΔM-Lip was less effective than wild-type lipin 1 in promoting expression of the CAAT enhancer binding protein α transcription factor (*Cebpa*), or mature adipocyte genes such as fatty acid binding protein 4 (*Fabp4*) and adiponectin (*Adipoq*) **(Fig. 5c)**. An independent replicate experiment also showed that ΔM-Lip lipin 1 was less effective at inducing lipogenic genes encoding acetyl-CoA carboxylase (*Acaca*) and diacylglycerol acyltransferase 1 (*Dgat1*) **(Supplementary Fig. 5c)**.

**Figure 5.**
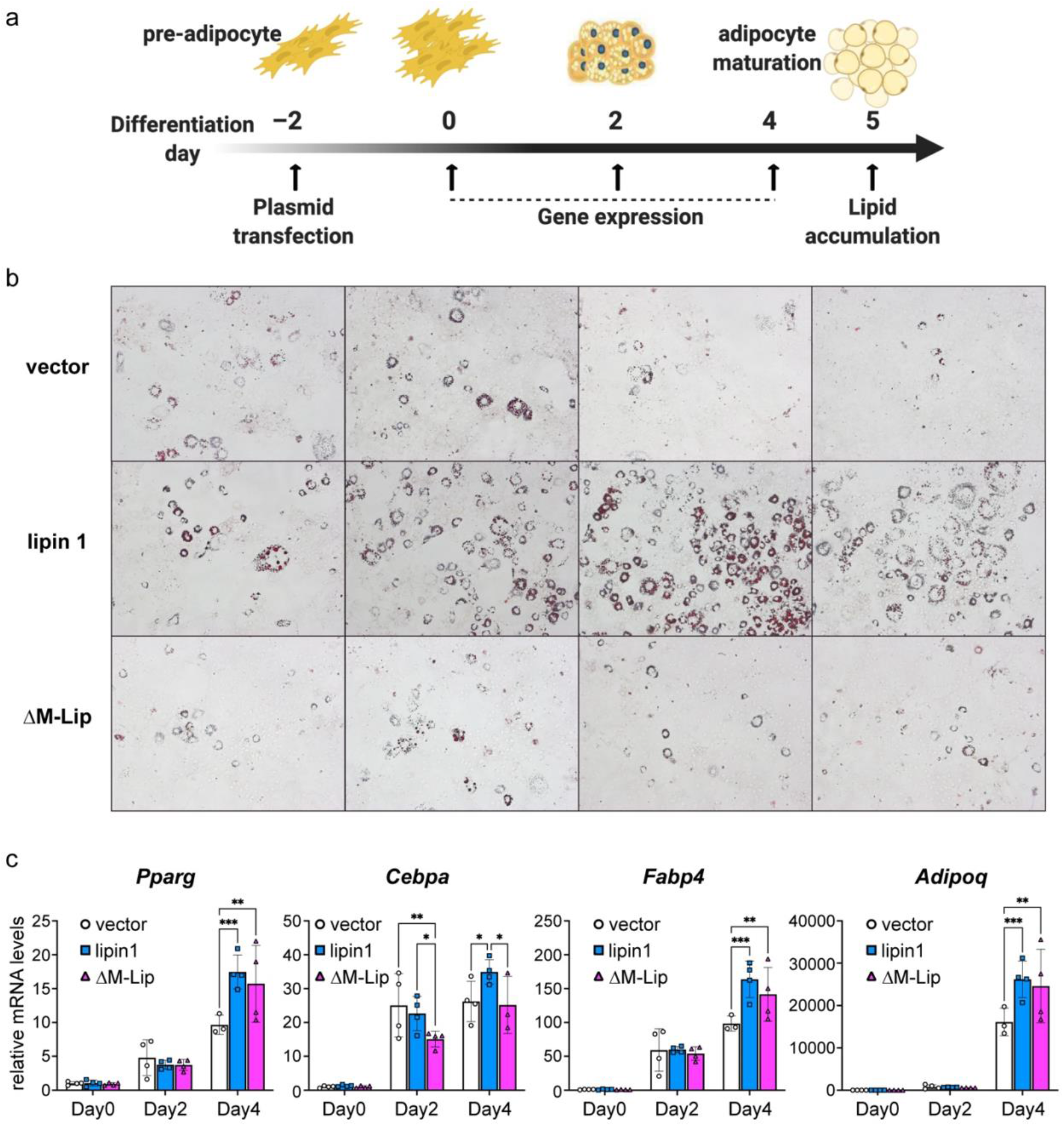
Optimal lipin 1 enhancement of adipogenesis requires the M-Lip region. **a**. The ability of wild-type and ΔM-Lip lipin 1 to promote 3T3-L1 preadipocyte differentiation were studied by transfection with corresponding expression constructs followed by analysis of gene expression and lipid accumulation. **b**. At day 5 after transfection, 3T3-L1 adipocytes were stained with oil red O to detect neutral lipid accumulation. Images shown are 100x magnification. **c**. Expression of genes encoding adipogenic transcription factors PPARγ (*Pparg*) and C/EBPα (*Cebpa*), adipocyte fatty acid binding protein FABP4 (*Fabp4*), and the adipocyte hormone, adiponectin (*Adipoq*), at days 0, 2, and 4 of differentiation. Gene expression was analyzed by 2-way ANOVA. *, p<0.05; **, p<0.01; ***, p<0.001. n=4 for each group.

## Discussion

This study identifies the M-Lip domain as a new protein fold that dimerizes and binds membranes. The M-Lip domain is functionally important for lipin subcellular localization, and enhancing adipogenesis, and can directly affect PAP activity in vitro in a manner that is dependent on membrane lipid content. With the exception of lipin 1 transcriptional co-activator function, deletion of the M-Lip had a negative effect but did not completely abrogate any lipin function. This is consistent with the sufficiency of the N-Lip and C-Lip regions for lipin PAP activity^18^ and all known disease mutations residing within the N-Lip or C-Lip regions.

We could identify the M-Lip domain in lipin homologs from several evolutionarily distant organisms, but not in *Sc* Pah1, nor in any non-lipin proteins from any species. Thus, the M-Lip domain appears to represent a unique feature of lipins and an evolutionary branch point that differentiates lipin PAP enzymes from mammals, invertebrates, ciliates, and plants from *Sc* Pah1.

Membrane association is the main regulatory mechanism that controls lipin PAP activity. Our HDX-MS studies reveal that membrane binding involves regions distributed throughout lipin 1. We propose mammalian lipins associate with membranes through a series of multi-valent interactions, with the M-Lip region contributing one site for membrane binding, as well as simultaneously doubling the number of membrane binding interactions through dimerization **(Fig. 6)**.

**Figure 6.**
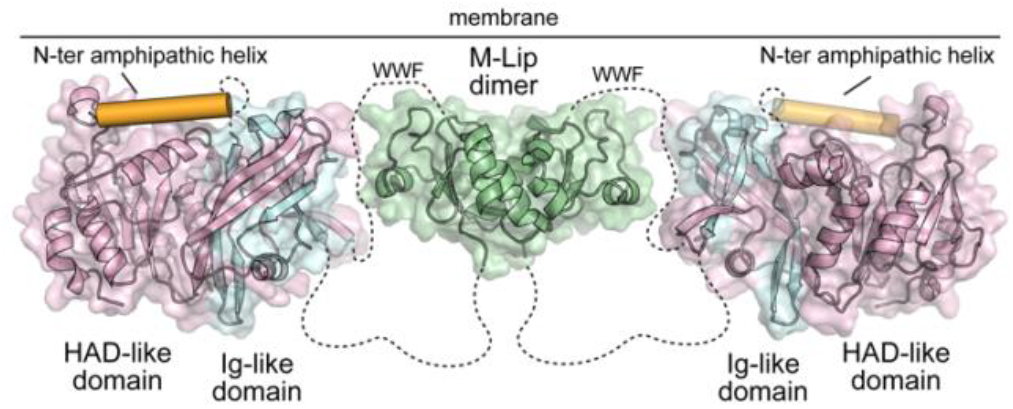
Structural model of mammalian lipin PAPs. Proposed model for a dimeric lipin 1 protein with the N-Lip (blue) and C-Lip (pink) combining to form the Ig-like and HAD-like domains. Membrane association is driven by multi-valent interactions from the N-terminal amphipathic helix (orange), the active site of the HAD-like domain, regions of the Ig-like domain, and the dimeric M-Lip that contains a hydrophobic WWF motif that is flanked by basic residues.

All of the individual membrane binding regions in lipins that have been characterized to date are responsive to the presence of anionic lipids, in particular PA^18, 22, 26, 27, 37^. This suggests that lipins may be recruited to cellular membranes in response to elevated PA levels. Given that PA is also the lipin enzyme substrate, this may reflect a mechanism to maintain low levels of PA.

As revealed by HDX-MS, the N-Lip and C-Lip regions are predominantly ordered in solution with substantial secondary structure. This finding suggests the N-Lip and C-Lip co-fold to form a split Ig-like domain and the HAD-like catalytic domain observed in *Tt* Pah2^18^. This is consistent with previous findings that a fusion of the N-Lip and C-Lip regions in both mouse lipin 2^18^ and *Sc* Pah1^32^ is sufficient for PAP activity in vitro. Notably, while the association of the N-Lip and C-Lip regions with one another appears to be constitutive in vitro, we cannot rule out that their association is transient or regulated in cells.

Here we expand the domain architecture of mammalian lipins to include M-Lip as a third protein domain that forms a stable dimer. Therefore, mammalian lipins must, at a minimum, be dimeric with the respective N-Lip and C-Lip regions co-folding either in cis (from the same subunit, **Fig. 6**) or in trans (from different subunits). We note that if the N-Lip and C-Lip interaction occurs in trans, this creates the potential to form larger oligomers when 4 or more lipin subunits combine. Lipin homo- and hetero-oligomerization has been demonstrated previously^34, 38^, and the size exclusion profile of recombinant lipin 1 yielded two peaks consistent with oligomerization beyond a dimer. Lastly, our data suggest that the conserved dimer interface of the M-Lip may mediate both homo and hetero-dimerization of mammalian lipins. This provides a foundation for more detailed experiments to unravel the functional consequences of lipin oligomerization in the regulation of phospholipid and triglyceride synthesis.

## Materials and Methods

### Plasmids

For expression in Sf9 insect cells, full-length mouse lipin 1 alpha (residues 1-891) was cloned into YM-Bac3 using SfoI and NotI restriction sites. YM-Bac3 is a modified version of pFastBac Htb (Invitrogen) that contains an N-terminal 6xHis tag followed by a Dual-Strep tag. The ΔM-Lip lipin 1 construct (residues 1-891 Δ458-565) was generated using the overlap extension method to delete the M-Lip domain and create an in-frame fusion between residues 457 and 566.

For E. coli expression, the mouse lipin 1 (M-Lip domain, residues 458-565; M-Lip^xtal^, residues 458-548) and mouse lipin 2 (M-Lip^xtal^, residues 459-549) were cloned into pET28b using NdeI and NotI restriction sites. Mouse lipin 1 M-Lip^xtal^ was also cloned into ppSUMO with BamHI and NotI sites, which added a ULP1 cleavable N-terminal His-tagged SUMO fusion.

For expression in mammalian cells, full-length mouse lipin 1α constructs (WT, residues 1-891; ΔM-Lip, residues 1-891 Δ458-565; and ΔM-Lip^xtal^, residues 1-891 Δ458-548) were cloned into pcDNA3.1 using BamHI and NotI restriction sites in frame with a C-terminal V5-His tag. ΔM-Lip, and ΔM-Lip^xtal^ deletion constructs were generated using the overlap extension method. The mouse lipin 1 M-Lip (mEGFP M-Lip) and M-Lip^xtal^ (mEGFP M-Lip^xtal^) domain fusions with monomeric enhanced GFP (mEGFP) located at the N-terminus were gene synthesized (BioBasic, Canada) and inserted into pcDNA3.1. Full-length mouse lipin 1 and the lipin 1 ΔM-Lip fusion with mEGFP located the C-terminus were generated by PCR and inserted into pcDNA3.1. All plasmids were directly sequenced.

### Protein expression and purification

#### Full-length mouse lipin 1 and ΔM-Lip lipin 1 proteins

Full-length mouse lipin 1 and ΔM-Lip lipin 1 were expressed in Sf9 cells using baculovirus. 300 mL of cells were infected with 1.0 mL of baculovirus at 3 million cells/mL at >95% viability and harvested 72 hours later. Cell pellets were lysed in 40 mM Tris, pH 8.0, 300 mM NaCl, 10 mM 2-mercaptoethanol by sonication, and the lysates were centrifuged at 26,000 rpm for 30 min. Lipin 1 and lipin 1 ΔM-Lip proteins were purified using Ni-NTA resin (GoldBio). Eluted protein was applied to Streptactin-XT resin equilibrated with equilibration buffer (100 mM Tris, pH 8.0, 150 mM NaCl, 10mM 2-mercaptoethanol), then washed with equilibration buffer, and eluted with equilibration buffer containing 50 mM biotin. Proteins were further purified by size exclusion chromatography using a Superdex 200 10/300 column in 20 mM Tris, pH 8.0, 150 mM NaCl, and 10 mM 2-mercaptoethanol. Fractions containing lipin 1 and ΔM-Lip lipin 1 proteins were concentrated, flash-frozen, and stored at −80°C.

#### Mouse lipin 1 M-Lip domain

The mouse lipin 1 M-Lip domain plasmid in pET28a was expressed in *E. coli* BL21 (DE3) RIPL cells. Cells were grown at 37 °C in Ultra-High Yield Flasks (Thompson Instrument Company) to an OD_600nm_ of 1.5 and then cooled at 10 °C for 2 hours. Protein expression was induced with isopropyl β-D-1-thiogalactopyranoside (IPTG) at 15 °C for 12 hrs, cells were harvested by centrifugation and stored at −80°C. Frozen cells were resuspended in buffer A (50 mM Tris-HCl, 60 mM imidazole, 500 mM NaCl, 5%(v/v) glycerol, 1% v/v Triton X-100, pH 7.4), lysed by sonication, centrifuged at 22,000 RPM (∼58,000 x g) at 4 °C for 1 hr, and applied to a gravity column with pre-equilibrated Ni-NTA resin (GoldBio). The column was washed with buffer A without Triton X-100, and protein was eluted with buffer B (50 mM Tris-HCl, 300 mM imidazole, 500 mM NaCl, 5% (v/v) glycerol, pH 7.4). The protein was diluted 4 fold in buffer C (50 mM HEPES, 50 mM NaCl, pH 7.35), applied to a HiTrap SP HP cation exchange column, washed with 5 column volumes of buffer C, and eluted with a linear gradient with buffer C supplemented with 1 M NaCl. Protein was aliquoted, flash frozen, and stored at −80 °C.

#### Mouse lipin 1 and mouse lipin 2 M-Lip^xtal^ domains

The M-Lip^xtal^ domains of mouse lipin 1 and mouse lipin 2 in pET28a were overexpressed in E. coli BL21 (DE3) RIPL cells. Cells were grown at 37 °C in Ultra-High Yield Flasks (Thompson Instrument Company) to an OD_600nm_ of 1.5 and then cooled at 10 °C for 2 hours. Protein expression was induced with isopropyl β-D-1-thiogalactopyranoside (IPTG) at 15°C for 18 hrs, cells were harvested by centrifugation, and stored at −80°C. Frozen cells were resuspended in buffer A (50 mM Tris-HCl, 60 mM imidazole, 500 mM NaCl, 5%(v/v) glycerol, pH 7.4), lysed by sonication, centrifuged at 22,000 RPM (∼58,000 x g) at 4 °C for 1 hr, and applied to a gravity column with pre-equilibrated Ni-NTA resin (GoldBio). The column was washed with buffer A and protein was eluted with buffer B (50 mM Tris-HCl, 300 mM imidazole, 500 mM NaCl, 5% (v/v) glycerol, pH 7.4). The eluted protein was applied to a HiLoad Superdex 75 26/600 column (GE Healthcare) equilibrated with SEC Buffer (50 mM Tris-HCl, 150 mM NaCl, 5% (v/v) glycerol, pH 7.4). Pooled fractions from SEC were concentrated to 10-20 mg/ml, aliquoted, flash frozen, and stored at −80°C.

#### Mouse lipin 1 M-Lip^xtal^ domain in ppSUMO

The mouse lipin 1 M-Lip^xtal^ domain that was crystallized was expressed and purified from the construct cloned into the ppSUMO plasmid. Expression conditions and the Ni-NTA purification protocol were identical as that described here for the M-Lip^xtal^ domain in pET28a. However, after elution from the Ni-NTA column, the protein was digested with purified ULP-1 overnight at 4°C. The digestion mixture was diluted 10 fold in buffer A, and re-applied to a Ni-column to remove the His-SUMO fusion. After, the M-Lip^xtal^ protein was further purified by SEC using a HiLoad Superdex 75 26/600 column (GE Healthcare) equilibrated with SEC Buffer. Pooled fractions from SEC were concentrated to 10-20 mg/ml, aliquoted, flash frozen, and stored at −80°C.

#### Se-Met derivatized mouse lipin 1 M-Lip^xtal^ protein

The mouse lipin 1 M-Lip^xtal^ domain derivatized with selenomethionine (Se-Met) was produced using B834 (DE3) cells. B834 cells were grown in M9 minimal medium supplemented with amino acids, Se-Met, and Kao and Michayluk Vitamin Solution (Sigma). Purification was identical to the M-Lip^xtal^ domain.

### Liposome generation

Large unilamellar vesicle (LUV) liposomes were prepared by the lipid extrusion method. Briefly, lipids dissolved in chloroform were dried under nitrogen gas for 30 minutes. For assays with full-length lipin 1 proteins, the liposomes were resuspended in 50 mM Tris, pH 7.5, 100 mM NaCl, 10 mM 2-mercaptoethanol buffer and LUVs were generated by 7 freeze-thaw cycles. Liposomes were composed of 20 mol% POPA (1-palmitoyl-2-oleoyl-sn-glycero-3-phosphate, Avanti Polar Lipids, #840857) and 0 or 40 mol% POPE (1-palmitoyl-2-oleoyl-sn-glycero-3-phosphoethanolamine, Avanti Polar Lipids, #850757) with the remaining 80 or 50 mol% as POPC (1-palmitoyl-2-oleoyl-glycero-3-phosphocholine, Avanti Polar Lipids, #850457).

For assays with the isolated M-Lip domains, liposomes were resuspended in PBS buffer. Followed by 3 rounds of freeze-thaw cycles with agitation, and extrusion with 100 nm size polycarbonate membrane filters (Avestin). Liposomes were composed of 73 mol% POPC, 5 mol% cholesterol (Avanti Polar Lipids, #700000P), 2 mol% PECF (1,2-dioleoyl-sn-glycero-3-phosphoethanolamine-N-(carboxyfluorescein), Avanti Polar Lipids, #810332) as visual guide and 20 mol% lipid of interest: POPA, POPS (1-palmitoyl-2-oleoyl-sn-glycero-3-phospho-L-serine, Avanti Polar Lipids, #840034), PODAG (1-palmitoyl-2-oleoyl-sn-glycerol, Avanti Polar Lipids, #800815), POPI (1-palmitoyl-2-oleoyl-sn-glycero-3-phosphoinositol, Avanti Polar Lipids, #850142) or ceramide (Avanti Polar Lipids, #8601518).

### Liposome sedimentation assays

For the full-length lipin 1 and ΔM-Lip lipin 1 proteins, 20 µL of LUV liposomes in 50 mM Tris, pH 7.5, 100 mM NaCl, 10 mM 2-mercaptoethanol buffer were mixed with 20 µL of proteins to give a final concentration of 1.0 mM liposomes and 1.0 µM protein. For the M-Lip and M-Lip^xtal^ proteins, 50 μL of LUV liposomes in pH 7.4 PBS buffer were mixed with 50 µL of proteins in PBS buffer, giving a final concentration of 1 mM liposomes and 50 µM protein. Reaction mixtures were incubated for 30 minutes and centrifuged at 100,000g at 4°C for 1 hr using a TLA100 fixed angle rotor (Beckman). The supernatant fraction was carefully removed, and the protein content of the pellet and supernatant fractions were analyzed by SDS-PAGE. All binding assays were performed at least three times and SDS-PAGE gel bands were quantified using ImageJ^39^.

### HDX-MS to determine ordered and disordered regions of full length lipin

HDX reactions were conducted in 20 µL reaction volumes with a final concentration of 0.63 µM full length lipin 1 per sample. Exchange was carried out in triplicate for a single time point (3 s at 4 °C). Hydrogen deuterium exchange was initiated by the addition of 18 µL of D_2_O buffer solution (20 mM HEPES pH 7, 100 mM NaCl, 0.5 mM TCEP) to the protein solution, to give a final concentration of 84.8% D_2_O. Exchange was terminated by the addition of acidic quench buffer at a final concentration 0.6 M guanidine-HCl and 0.9% formic acid. Samples were immediately frozen in liquid nitrogen at −80 °C. Fully deuterated samples were generated by first denaturing the protein in 6M guanidine for 1 hour at RT. Following denaturing, 18 μL of D_2_O buffer was added to the denatured protein and allowed to incubate for 15 minutes at RT before quenching. The 3s on ice (1 °C) condition was created and run in triplicate. Samples were flash frozen in liquid nitrogen and stored at −80°C until injection onto an ultra-performance liquid chromatography system for proteolytic cleavage, peptide separation, and injection onto a QTOF for mass analysis.

### HDX-MS mapping of full-length lipin 1 with liposomes

Liposomes were made by resuspending the lipid film (20% POPA, 80% POPC) in lipid buffer (50 mM Tris pH 7.5, 0.1 M NaCl, 10 mM βME) resulting in a final liposome concentration of 2 mM. The resuspended lipid was sonicated for 10 minutes, freeze-thawed seven times, and then extruded through a prewet 100nm filter unit 10x times prior to usage in the experiment. HDX reactions were conducted in a final reaction volume of 20 μL with a lipin 1 protein concentration of 0.63 μM. Prior to the addition of deuterated solvent, 1 μL of lipin 1 was allowed to incubate with either 2 μL of 2 mM liposomes or 2 μL of the corresponding lipid buffer. After the two-minute incubation period, 17 μL D_2_O buffer (20mM HEPES pH 7, 100mM NaCl, 0.5 mM TCEP, 94% D_2_O) was added with a final %D_2_O of 80.1% (v/v). The reaction was allowed to proceed for 3s, 30s, 300s, or 3000s at 18°C before being quenched with ice cold acidic quench buffer. All conditions and timepoints were created and run in triplicate. Samples were flash frozen in liquid nitrogen and stored at −80°C until injection onto an ultra-performance liquid chromatography system for proteolytic cleavage, peptide separation, and injection onto a QTOF for mass analysis.

### HDX-MS mapping of the M-Lip domain with liposomes

Liposomes were made by resuspending the lipid film (20% POPA, 60% POPC, 20% POPE) in lipid buffer (20 mM HEPES pH 7.5, 100 mM NaCl) resulting in a final liposome concentration of 4 mM. The resuspended lipid was sonicated for 10 minutes, freeze-thawed three times, and then extruded through a prewet 100nm filter unit 11x times prior to usage in the experiment. HDX reactions were conducted in a final reaction volume of 20 µL with an M-Lip concentration of 1.2 µM. Prior to the addition of deuterated solvent, 1 µL M-Lip protein was allowed to incubate with either 1 µL of 4 mM lipid vesicles or 1 µL of the corresponding lipid buffer. After a two minute incubation period, 18 µL D_2_O buffer (20 mM HEPES pH 7.5, 100 mM NaCl, 94% D_2_O (v/v)) was added with a final %D_2_O of 84.9% (v/v). The reaction allowed to proceed for 3s, 30s, 300s, or 3000s at 18°C before being quenched with ice cold acidic quench buffer, resulting in a final concentration of 0.6 M guanidine-HCl and 0.9% FA post quench. All conditions and timepoints were created and run in triplicate. Samples were flash frozen in liquid nitrogen and stored at −80°C until injection onto an ultra-performance liquid chromatography system for proteolytic cleavage, peptide separation, and injection onto a QTOF for mass analysis.

### Protein digestion and MS/MS data collection

Protein samples were rapidly thawed and injected onto an integrated fluidics system containing a HDx-3 PAL liquid handling robot and climate-controlled chromatography system (LEAP Technologies), a Dionex Ultimate 3000 UHPLC system, as well as an Impact HD QTOF Mass spectrometer (Bruker). The protein was run over either one (at 10°C) or two (at 10°C and 2°C) immobilized pepsin columns (Applied Biosystems; Poroszyme Immobilized Pepsin Cartridge, 2.1 mm x 30 mm; Thermo-Fisher 2-3131-00; Trajan; ProDx protease column, 2.1 mm x 30 mm PDX.PP01-F32) at 200 mL/min for 3 minutes. The resulting peptides were collected and desalted on a C18 trap column (Acquity UPLC BEH C18 1.7mm column (2.1 x 5 mm); Waters 186003975). The trap was subsequently eluted in line with a C18 reverse −phase separation column (Acquity 1.7 mm particle, 100 x 1 mm^2^ C18 UPLC column, Waters 186002352), using a gradient of 5-36% B (Buffer A 0.1% formic acid; Buffer B 100% acetonitrile) over 16 minutes. Mass spectrometry experiments acquired over a mass range from 150 to 2200 m/z using an electrospray ionization source operated at a temperature of 200C and a spray voltage of 4.5 kV.

### Peptide identification

Peptides were identified using data-dependent acquisition following tandem MS/MS experiments (0.5 s precursor scan from 150-2000 m/z; twelve 0.25 s fragment scans from 150-2000 m/z). MS/MS datasets were analyzed using PEAKS7 (PEAKS), and peptide identification was carried out by using a false discovery-based approach, with a threshold set to 1% using a database of purified proteins and known contaminants. The search parameters were set with a precursor tolerance of 20 ppm, fragment mass error 0.02 Da, and charge states from 1-8.

### Mass analysis of peptide centroids and measurement of deuterium incorporation

HDExaminer Software (Sierra Analytics) was used to automatically calculate the level of deuterium incorporation into each peptide. All peptides were manually inspected for correct charge state and presence of overlapping peptides. Deuteration levels were calculated using the centroid of the experimental isotope clusters. The results for the experiment comparing lipin with and without liposomes are presented as relative levels of deuterium incorporation, with no control for back exchange, and the only correction was for the level of deuterium present in the buffer (either 84.9% or 80.1%). The fully deuterated sample in the order-disorder experiment allowed for a back-exchange correction in this specific experiment during digestion and separation. Differences in exchange in a peptide were considered significant if they met all three of the following criteria: >5% change in exchange, >0.4 Da difference in exchange, and a *p*-value <0.01 using a two-tailed Student’s *t*-test. All compared samples were set within the same experiment. The raw HDX data are shown in two different formats. The raw peptide deuterium incorporation graphs for a selection of peptides with significant differences are shown, with the raw data for all analyzed peptides in the source data. To allow for visualization of differences across all peptides, we utilized number of deuteron difference (#D) plots. These plots show the total difference in deuterium incorporation over the entire H/D exchange time course, with each point indicating a single peptide. The data analysis statistics for all HDX-MS experiments are in Supplemental Table 1 according to the guidelines of Masson et al^29^. The mass spectrometry proteomics data have been deposited to the ProteomeXchange Consortium via the PRIDE partner repository^40^ with the dataset identifier PXD022172.

### Crystallization and data collection

All crystals were grown using the hanging-drop method by mixing 1.5 μL of reservoir solution mixed with 1.5 μL of protein solution at RT. Crystals of native and Se-Met mouse lipin 1 M-Lip^xtal^ domain were grown in 5-15% PEG 3000, 0.1 M Tris-HCl, pH 7.0 by seeding with microcrystals generated using a Seed Bead Kit (Hampton Research, HR4-781). Crystal form 2 of mouse lipin 1 M-Lip^xtal^ were grown in 0.2 M Zinc Acetate, 1 M NaCl, 0.1 M Imidazole, pH 8.0 by seeding with microcrystals. Crystals of mouse lipin 2 M-Lip^xtal^ were grown in 0.2 M Calcium Acetate, 20% PEG 3350. All crystals were cryoprotected with 25% v/v glycerol and flash frozen in liquid nitrogen prior to data collection. Diffraction data for mouse lipin 1 native M-Lip^xtal^ domain was collected at 0.979 Å at the APS NE-CAT 24-ID-C beamline **(Table1)**. Data for the Se-Met and crystal form 2 of the M-Lip^xtal^ domain were collected at 0.979Å and 1.28Å at the NSLS-II FMX 17-ID beamline at Brookhaven National Laboratory **(Table 1)**. Diffraction data for mouse lipin 2 M-Lip^xtal^ was collected at 0.979Å at the Advanced Photon Source GM/CA ID-23B beamline at Argonne National Laboratory **(Table 1)**.

### Data processing, structure determination, and refinement

Diffraction data was integrated and scaled using XDS-DIALS^41^ in CCP4^42^. Se-Met SAD phases were calculated to 2.3Å resolution using Phenix AutoSol^43^, which generated an initial map. The resulting map showed clear electron density for side-chains and the model was further improved by manual model building using COOT^44^. This yielded a nearly complete search model for molecular replacement with the 1.5 Å resolution native data set. Subsequent model adjustments were carried out in COOT with further refinement with Phenix^45^. The finalized structure mouse lipin 1 M-Lip^xtal^ domain was used as a search model for molecular replacement for crystal form 2 of mouse lipin 1 and the mouse lipin 2 M-Lip^xtal^ domain using Phaser^46^ in Phenix^45^. The final models were generated by manual model building in COOT and refinement in Phenix. Data collection and refinement statistics are provided in Table 1.

### SEC-MALS

1.6 mg of the mouse lipin 1 M-Lip^xtal^ domain was injected onto a Superdex 200 Increase 10/300 (GE health) size exclusion column with constant flow rate of 0.35 ml/min. Light scattering and refractive index data were collected by miniDAWN TREOS (Wyatt technology) and Optiab T-rEX (Wyatt technology), respectively. The refractive index change was measured differentially by miniDAWN TREOS with a laser at a wavelength of 658 nm, and UV absorbance was measured with the diode array detector at 280 nm. Data were subsequently processed by ASTRA software 7 (Wyatt technology) and peak alignment and band broadening correction between the UV, MALS, and RI detectors were performed using Astra software algorithms.

### Cell culture and transfection

Murine Hepa1-6 hepatoma cells (American Type Culture Collection #CRL-1830, Manassas, VA) and human HEK293 embryonic kidney cells (CRL-1573) were maintained in minimal essential medium (MEM) with 10% fetal bovine serum (FBS) (Corning Inc., Corning, NY). All experiments were performed with cells having 2–6 passages. Cells were transfected with plasmids using BioT reagent (Bioland Scientific, Paramount, CA).

### Co-Immunoprecipitation

Co-immunoprecipitation experiments were performed to detect lipin 1 dimer formation. To unambiguously identify the interaction between two lipin monomers, Hepa1-6 cells were transfected with independent lipin 1 constructs that were fused to different epitopes (V5 or BirA*-HA). To detect lipin1 heterodimer formation, lipin 1 constructs with V5 epitope (wild-type lipin 1-V5, ΔMLip-V5, or ΔM-Lip^xtal^-V5) were co-transfected with lipin 1-BirA*-HA, lipin 2-BirA*-HA or lipin 3-BirA*-HA. Two days after transfection, cells were harvested and lysed in 0.1% NP-40 with phosphatase inhibitor cocktails (Sigma-Aldrich, St. Louis, MO). After brief sonication, the supernatants were collected by centrifugation at 12,000 rpm for 10 min at 4 °C. For immunoprecipitation of V5-tagged proteins, cell lysates were incubated with anti-V5 antibody (ThermoFisher Scientific, R960-25, Waltham, MA) at 4°C overnight. Cell lysate/antibody mixture were then incubated with protein A/G-agarose beads (Santa Cruz Biotechnology, Inc., Santa Cruz, CA) for 2-hour at 4°C. The precipitates were washed three times with lysis buffer, boiled in sample buffer for 5 min and subjected to immunoblot assay with anti-HA antibody (Cell Signaling Technology, Inc., #3724, Danvers, MA).

### Lipin 1 transcriptional coactivation assay

Ability of wild-type and ΔM-Lip lipin 1 to coactivate PGC-1α was assessed as described previously^8, 15^. Briefly, HEK293 cells were transfected with a PPRE-firefly luciferase reporter plasmid together with phRL-TK *Renilla* luciferase control vector (Promega, Madison, WI), pCMX-PPARα, pCMX-PGC-1α, pCMX-RXR, and wild-type lipin 1, ΔMLip-V5, or lipin 1 ΔM-Lip^xtal^-V5 expression constructs. Two days after transfection, cells were collected and luciferase activity determined with the Dual-Luciferase Assay System (Promega, Madison, WI).

### 3T3-L1 adipogenesis

3T3-L1 preadipocytes were cultured and differentiated as described previously^47^. To investigate the function of the M-Lip domain in adipogenesis, cells were reverse-transfected at differentiation day –2 (confluent) with pcDNA, wild-type lipin 1-V5 or ΔM-Lip-V5 (BioT reagent, Bioland Scientific). At day 0, cells were changed to differentiation medium (ZenBio, DM2L1500), and at day 2 were switched to maintenance medium (ZenBio, AM-1-L1). RNA isolation was performed at day 0, day 2 and day 4 of differentiation. Cell morphology was monitored by brightfield phase-contrast and Oil red O staining.

### RNA extraction and quantitative RT-PCR

Total RNA was isolated from 3T3-L1 cells using TRIzol (Life Technologies) and reverse transcribed (iScript, Bio-Rad). Real-time RT-PCR was performed with a Bio-Rad CFX Connect Real-Time PCR Detection System using SsoAdvanced SYBR Green Supermix and *Tbp* mRNA as a normalization control. Primers were as follows:

*Pparg* (AACTCTGGGAGATTCTCCTGTTGA, TGGTAATTTCTTGTGAAGTGCTCATA);

*Cebpa* (GAACAGCAACGAGTAACCGGGTA, GCCATGGCCTTGACCAAGGAG);

*Fabp4* (AACCTGGAAGCTTGTCTCCA, CACGCCCAGTTTGAAGGAAA);

*Adipoq* (TGTTCCTCTTAATCCTGCCCA, CCAACCTGCACAAGTTCCCTT);

*Acaca* (GCCTCTTCCTGACAAACGAG, TGACTGCCGAAACATCTCTG);

*Fasn* (GTTGGCCCAGAACTCCTGTA, GTCGTCTGCCTCCAGAGC);

*Dgat1* (CTGAATTGGTGTGTGGTGATG, AGGGGTCCTTCAGAAACAGAG);

*Tbp* (ACCCTTCACCAATGACTCCTATG, ATGATGACTGCAGCAAATCGC).

### Oil red O stain

At day 5 of adipocyte differentiation, cells were fixed in 10% formalin for 20 min, washed with 60% isopropanol, and stained with oil red O solution (0.2% w/v in isopropanol). Cell images were captured at 100x magnification. Pictures from 4 independent wells per treatment group were used.

### PAP activity assays

Micelles containing 5 mol% nitrobenzoxadiazole-phosphatidic acid (NBD-PA) (Avanti Polar Lipids, #9000341) and Triton X-100 (Research Products International Corp, #400001) were generated in a buffer containing 50 mM Tris pH 7.5, 100 mM NaCl, 10 mM 2-mercaptoethanol, and 4 mM MgCl_2_, to give a final bulk concentration of 80 µM NBD-PA. Liposomes used for PAP assays were composed of 10 mol% NBD-PA with 0 mol% or 40% POPE with the remaining as POPC (90 mol% or 50 mol%) and prepared in buffer containing 50 mM Tris, pH 7.5, 100 mM NaCl, 10 mM 2-mercaptoethanol, and 4 mM MgCl_2_ to give a final bulk concentration of 150 µM NBD-PA.

In a 50 µL reaction mixture, 45 µL NBD-PA mixed micelles or liposomes were incubated with 5 µL lipin 1 or lipin 1 ΔM-Lip (total 5 ng protein) for 30 min at 30 °C. Reactions were quenched by addition of 150 µL of CHCl_3_/MeOH (1:1), then vortexed and centrifuged at 2,000 rpm for 3 min. The organic phase was collected, dried under N_2_(g), and resuspended with 100 µL methanol containing 0.2% formic acid and 1 mM ammonium formate.

Both the mixed-micelle and liposome samples were analyzed by HPLC using a Spectra 3µm C8SR column (3 µm particle, 150 x 3.0 mm, Peeke Scientific, #S-3C8SR-FJ) under the following conditions: solvent A: water containing 0.2% formic acid and 1 mM ammonium formate; solvent B: methanol containing 0.2% formic acid and 1 mM ammonium formate; flow rate: 0.5ml/min. The gradient profile started at 80% for solvent B and was increased to 98% B after 7 min, then kept at 98% for 3 min. 10 µL samples were injected onto the column, which was kept at 35 °C at all runs. The fluorescent signal was detected at excitation and <u>emission wavelengths</u>of 470 and 530 nm, respectively. The detector signal was recorded and integrated by a using Agilent technology OpenLAB CDS ChemStation edition software.

### Confocal microscopy

COS-7 cells (American Type Culture Collection, CRL-1651) were seeded in 6-well plate with a cover glass inside and were transfected using lipofectamine 2000 (Thermo Fisher, #11668027) with mouse mEGFP M-Lip or mEGFP M-Lip^xtal^ and mApple-Sec61b, or lipin 1 mEGFP or ΔM-Lip lipin 1 mEGFP in pcDNA3.1. 48 hr after transfection cells were rinsed with PBS three times, then fixed with 4% paraformaldehyde containing 5 µg/ml Hoechst 33342 stain (Thermo Fisher, #62249) for 10 min at room temperature, then rinsed three times with PBS. The cover glass was then mounted onto a microslide. The cells were visualized using Zeiss Axio imager M2 confocal microscope.

## Data availability

Coordinates and structure factors have been deposited in the Protein Data Bank under accession codes 7KIH, 7KIL, 7KIQ. The mass spectrometry proteomics data have been deposited to the ProteomeXchange Consortium via the PRIDE partner repository^40^ with the dataset identifier PXD022172. All other data are available from the authors on request. The source data underlying Figs 1b-e, 2e-f, 3a, 3c-f, 4a-c, 5c, and supplementary figure 5c are provided as a Source Data file.

## Acknowledgements

We thank the staff at the NSLS-II (AMX and FMX) and APS (GM-CAT and NE-CAT) beamlines for assistance during data collection, and Vitaly Citovsky and Benoit Lacroix (Stony Brook) for access to their confocal microscope supported by the NIH grant 5R01GM05022423. This work was supported by the National Institutes of Health grants R35 GM128666 (MVA), P01 HL090553 (KR), and P01 HL028481 (KR), the American Heart Association grants 17SDG33410860 (MVA) and 18POST34060200 (HW), the NSERC Discovery Grant NSERC-2020-04241 (JEB), the Michael Smith Foundation for Health Research (JEB, Scholar Award 17686), and a Stony Brook URECA award (NMP).

## Author contributions

WG performed all experiments with the isolated mouse lipin 1 M-Lip domain including protein purifications, crystallization experiments, SEC-MALS, and liposome sedimentation assays. JWY purified and crystallized the mouse lipin 2 M-Lip^xtal^ domain. WG and MVA determined and refined the final crystal structures. SG purified full-length lipin 1 and ΔM-Lip proteins and performed PAP assays, liposome sedimentation assays, and confocal microscopy experiments. KDF and RMH performed the HDX-MS experiments for lipin 1 and the M-Lip, respectively. KDF, RMH, and JEB analyzed all HDX-MS data. HW performed co-immunoprecipitation, transcriptional co-activator, and adipogenesis studies. YMC conceived of and generated the YM-Bac3 plasmid. NMP generated key constructs for experiments. WG, SG, HW, KR, JEB, and MVA contributed intellectual and strategic input. KR, JEB, and MVA supervised work and provided funding support. WG, SG, KR, and MVA drafted the initial manuscript with contributions from KDF, JEB, and HW. WG, SG, KR, JEB, KR, and MVA edited the final manuscript. All authors approved the final manuscript.

## Competing Interest Statement

The authors have no conflicting interest to report.

## Notes

### Competing Interest Statement

The authors have declared no competing interest.

## References

1. Han G-S, Wu W-I, Carman GM. The Saccharomyces cerevisiae Lipin homolog is a Mg2+-dependent phosphatidate phosphatase enzyme. Journal of Biological Chemistry 281, 9210–9218 (2006).

2. Siniossoglou S. Phospholipid metabolism and nuclear function: roles of the lipin family of phosphatidic acid phosphatases. Biochimica et Biophysica Acta (BBA)-Molecular and Cell Biology of Lipids 1831, 575–581 (2013).

3. Lutkewitte AJ, Finck BN. Regulation of Signaling and Metabolism by Lipin-mediated Phosphatidic Acid Phosphohydrolase Activity. Biomolecules 10, 1386 (2020).

4. Zhang P, et al. Lipin 2/3 phosphatidic acid phosphatases maintain phospholipid homeostasis to regulate chylomicron synthesis. The Journal of clinical investigation 129, 281–295 (2019).

5. Phan J, Péterfy M, Reue K. Lipin expression preceding peroxisome proliferator-activated receptor-γ is critical for adipogenesis in vivo and in vitro. Journal of Biological Chemistry 279, 29558–29564 (2004).

6. Péterfy M, Phan J, Xu P, Reue K. Lipodystrophy in the fld mouse results from mutation of a new gene encoding a nuclear protein, lipin. Nature genetics 27, 121–124 (2001).

7. Harris TE, et al. Insulin controls subcellular localization and multisite phosphorylation of the phosphatidic acid phosphatase, lipin 1. Journal of Biological Chemistry 282, 277–286 (2007).

8. Donkor J, et al. A conserved serine residue is required for the phosphatidate phosphatase activity but not the transcriptional coactivator functions of lipin-1 and lipin-2. Journal of Biological Chemistry 284, 29968–29978 (2009).

9. Schweitzer GG, et al. Rhabdomyolysis-associated mutations in human LPIN1 lead to loss of phosphatidic acid phosphohydrolase activity. In: JIMD Reports, Volume 23). Springer (2015).

10. Michot C, et al. LPIN1 gene mutations: a major cause of severe rhabdomyolysis in early childhood. Human mutation 31, E1564–E1573 (2010).

11. Ferguson P, et al. Homozygous mutations in LPIN2 are responsible for the syndrome of chronic recurrent multifocal osteomyelitis and congenital dyserythropoietic anaemia (Majeed syndrome). Journal of medical genetics 42, 551–557 (2005).

12. Zeharia A, et al. Mutations in LPIN1 cause recurrent acute myoglobinuria in childhood. The American Journal of Human Genetics 83, 489–494 (2008).

13. Zhang P, Verity MA, Reue K. Lipin-1 regulates autophagy clearance and intersects with statin drug effects in skeletal muscle. Cell metabolism 20, 267–279 (2014).

14. Yao-Borengasser A, et al. Lipin expression is attenuated in adipose tissue of insulin-resistant human subjects and increases with peroxisome proliferator–activated receptor γ activation. Diabetes 55, 2811–2818 (2006).

15. Finck BN, et al. Lipin 1 is an inducible amplifier of the hepatic PGC-1α/PPARα regulatory pathway. Cell metabolism 4, 199–210 (2006).

16. Donkor J, Sariahmetoglu M, Dewald J, Brindley DN, Reue K. Three mammalian lipins act as phosphatidate phosphatases with distinct tissue expression patterns. Journal of Biological Chemistry 282, 3450–3457 (2007).

17. Carman GM, Han G-S. Fat-regulating phosphatidic acid phosphatase: a review of its roles and regulation in lipid homeostasis. Journal of lipid research 60, 2–6 (2019).

18. Khayyo VI, et al. Crystal structure of a lipin/Pah phosphatidic acid phosphatase. Nature communications 11, 1–11 (2020).

19. Harris TE, Finck BN. Dual function lipin proteins and glycerolipid metabolism. Trends in Endocrinology & Metabolism 22, 226–233 (2011).

20. Huffman TA, Mothe-Satney I, Lawrence JC. Insulin-stimulated phosphorylation of lipin mediated by the mammalian target of rapamycin. Proceedings of the National Academy of Sciences 99, 1047–1052 (2002).

21. Péterfy M, Harris TE, Fujita N, Reue K. Insulin-stimulated interaction with 14-3-3 promotes cytoplasmic localization of lipin-1 in adipocytes. Journal of Biological Chemistry 285, 3857–3864 (2010).

22. Eaton JM, Mullins GR, Brindley DN, Harris TE. Phosphorylation of lipin 1 and charge on the phosphatidic acid head group control its phosphatidic acid phosphatase activity and membrane association. Journal of Biological Chemistry 288, 9933–9945 (2013).

23. Liu G-H, Gerace L. Sumoylation regulates nuclear localization of lipin-1α in neuronal cells. PloS one 4, e7031 (2009).

24. Li TY, et al. Tip60-mediated lipin 1 acetylation and ER translocation determine triacylglycerol synthesis rate. Nature communications 9, 1–14 (2018).

25. Peterson TR, et al. mTOR complex 1 regulates lipin 1 localization to control the SREBP pathway. Cell 146, 408–420 (2011).

26. Karanasios E, Han G-S, Xu Z, Carman GM, Siniossoglou S. A phosphorylation-regulated amphipathic helix controls the membrane translocation and function of the yeast phosphatidate phosphatase. Proceedings of the National Academy of Sciences 107, 17539–17544 (2010).

27. Ren H, et al. A phosphatidic acid binding/nuclear localization motif determines lipin1 function in lipid metabolism and adipogenesis. Molecular biology of the cell 21, 3171–3181 (2010).

28. Boroda S, et al. The phosphatidic acid–binding, polybasic domain is responsible for the differences in the phosphoregulation of lipins 1 and 3. Journal of Biological Chemistry 292, 20481–20493 (2017).

29. Masson GR, et al. Recommendations for performing, interpreting and reporting hydrogen deuterium exchange mass spectrometry (HDX-MS) experiments. Nature methods 16, 595–602 (2019).

30. Vadas O, Burke JE. Probing the dynamic regulation of peripheral membrane proteins using hydrogen deuterium exchange–MS (HDX–MS). Biochemical Society Transactions 43, 773–786 (2015).

31. Burke JE. Dynamic structural biology at the protein membrane interface. Journal of Biological Chemistry 294, 3872–3880 (2019).

32. Park Y, Han G-S, Carman GM. A conserved tryptophan within the WRDPLVDID domain of yeast Pah1 phosphatidate phosphatase is required for its in vivo function in lipid metabolism. Journal of Biological Chemistry 292, 19580–19589 (2017).

33. Holm L, Rosenstrom P. Dali server: conservation mapping in 3D. Nucleic acids research 38, W545–W549 (2010).

34. Liu G-H, et al. Lipin proteins form homo-and hetero-oligomers. Biochemical Journal 432, 65–76 (2010).

35. Zhang P, Takeuchi K, Csaki LS, Reue K. Lipin-1 phosphatidic phosphatase activity modulates phosphatidate levels to promote peroxisome proliferator-activated receptor γ (PPARγ) gene expression during adipogenesis. Journal of Biological Chemistry 287, 3485–3494 (2012).

36. Mitra MS, et al. Mice with an adipocyte-specific lipin 1 separation-of-function allele reveal unexpected roles for phosphatidic acid in metabolic regulation. Proceedings of the National Academy of Sciences 110, 642–647 (2013).

37. Eaton JM, et al. Lipin 2 binds phosphatidic acid by the electrostatic hydrogen bond switch mechanism independent of phosphorylation. Journal of Biological Chemistry 289, 18055–18066 (2014).

38. Creutz CE, Eaton JM, Harris TE. Assembly of high molecular weight complexes of lipin on a supported lipid bilayer observed by atomic force microscopy. Biochemistry 52, 5092–5102 (2013).

39. Rasband WS. ImageJ.). Bethesda, MD (1997).

40. Perez-Riverol Y, et al. The PRIDE database and related tools and resources in 2019: improving support for quantification data. Nucleic acids research 47, D442–D450 (2019).

41. Winter G, et al. DIALS: implementation and evaluation of a new integration package. Acta Crystallographica Section D 74, 85–97 (2018).

42. Potterton L, et al. CCP4i2: the new graphical user interface to the CCP4 program suite. Acta Crystallographica Section D: Structural Biology 74, 68–84 (2018).

43. Liebschner D, et al. Macromolecular structure determination using X-rays, neutrons and electrons: recent developments in Phenix. Acta Crystallographica Section D: Structural Biology 75, 861–877 (2019).

44. Emsley P, Cowtan K. Coot: model-building tools for molecular graphics. Acta Crystallographica Section D: Biological Crystallography 60, 2126–2132 (2004).

45. Adams PD, et al. PHENIX: a comprehensive Python-based system for macromolecular structure solution. Acta Crystallographica Section D: Biological Crystallography 66, 213–221 (2010).

46. McCoy AJ, Grosse-Kunstleve RW, Adams PD, Winn MD, Storoni LC, Read RJ. Phaser crystallographic software. Journal of applied crystallography 40, 658–674 (2007).

47. Link JC, et al. X chromosome dosage of histone demethylase KDM5C determines sex differences in adiposity. The Journal of Clinical Investigation 130, (2020).

