## Supplementary Information for "The middle lipin (M-Lip) domain is a new dimeric protein fold that binds membranes"

Supplementary Information – Gu et al

### Supplementary Table 1. HDX Data Summary

| Data Set | lipin-1 order/disorder | lipin-1 | lipin-1 + membrane | M-Lip domain | M-Lip domain + membrane |
|---|---|---|---|---|---|
| HDX reaction details | $\%D_2O$=84.8% pH$_{(read)}$= 7.5 Temp= 18°C | $\%D_2O$=80.1% pH$_{(read)}$= 7.5 Temp= 18°C | $\%D_2O$=80.1% pH$_{(read)}$= 7.5 Temp= 18°C | $\%D_2O$=85% pH$_{(read)}$= 7.5 Temp= 18°C | $\%D_2O$=85% pH$_{(read)}$= 7.5 Temp= 18°C |
| HDX time course | 3s at 4°C | 3s, 30s, 300s, 3000s | 3s, 30s, 300s, 3000s | 3s, 30s, 300s, 3000s | 3s, 30s, 300s, 3000s |
| HDX controls | N/A | N/A | N/A | N/A | N/A |
| Back-exchange | Corrected based on $\%D_2O$ | Corrected based on $\%D_2O$ | Corrected based on $\%D_2O$ | Corrected based on $\%D_2O$ | Corrected based on $\%D_2O$ |
| Number of peptides | 189 | 179 | 179 | 82 | 82 |
| Sequence coverage | 88.8% | 90.6% | 90.6% | 99.2% | 99.2% |
| Average peptide length / Redundancy | Length = 15.8 Redundancy = 3.1 | Length = 15.9 Redundancy = 3.0 | Length = 15.9 Redundancy = 3.0 | Length = 13.2 Redundancy = 8.4 | Length = 13.2 Redundancy = 8.4 |
| Replicates | 3 | 3 | 3 | 3 | 3 |
| Repeatability | Average StDev = 0.6% | Average StDev = 0.8% | Average StDev = 0.7% | Average StDev = 0.5% | Average StDev = 0.5% |
| Significant differences in HDX | N/A | >4% and >0.4 Da and unpaired t-test <0.01 | >4% and >0.4 Da and unpaired t-test <0.01 | >5% and >0.4 Da and unpaired t-test <0.01 | >5% and >0.4 Da and unpaired t-test <0.01 |

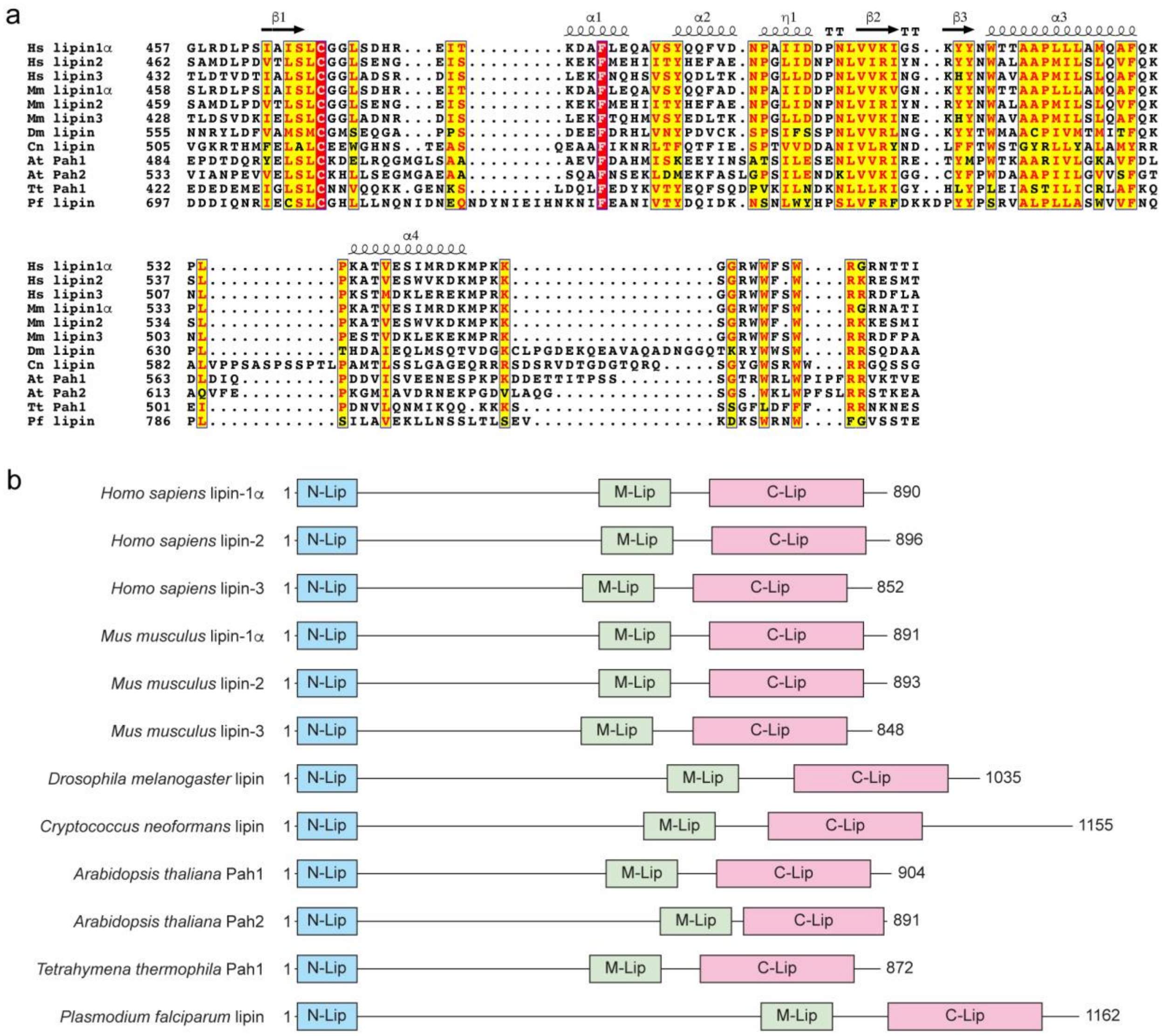

**Supplementary Figure 1. Evolutionary conservation of the M-Lip domain**

**(a)** Sequence alignment of the M-Lip domain from evolutionarily distant organisms. Identical residues are shaded red and residues with positive homology are shaded yellow. Secondary structure elements for the mouse lipin 1 M-Lip domain are indicated above, "TT" = turn.

**(b)** Domain architecture of lipin/Pahs that contain the M-Lip domain, drawn to scale. Other conserved motifs, e.g. nuclear localization signals, predicted amphipathic helices, and the conserved Trp motif are omitted for clarity.

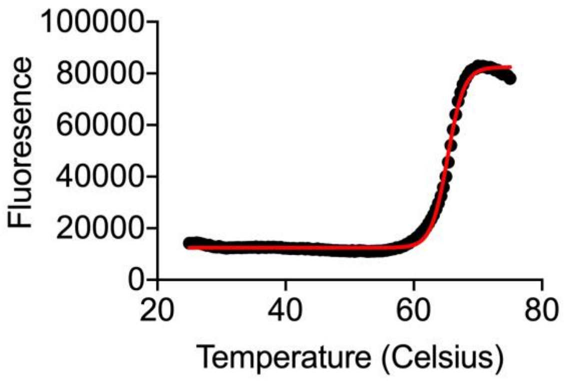


**Supplementary Figure 2.** The thermal stability of the mouse lipin-1 M-Lip$^{xtal}$ domain was assessed by differential scanning fluorimetry using Sypro Orange (Sigma Aldrich). Individual data points are shown as black circles. A Boltzmann sigmoidal fit (red line) was used to determine the melting temperature. The data indicate a melting temperature of 65°C.

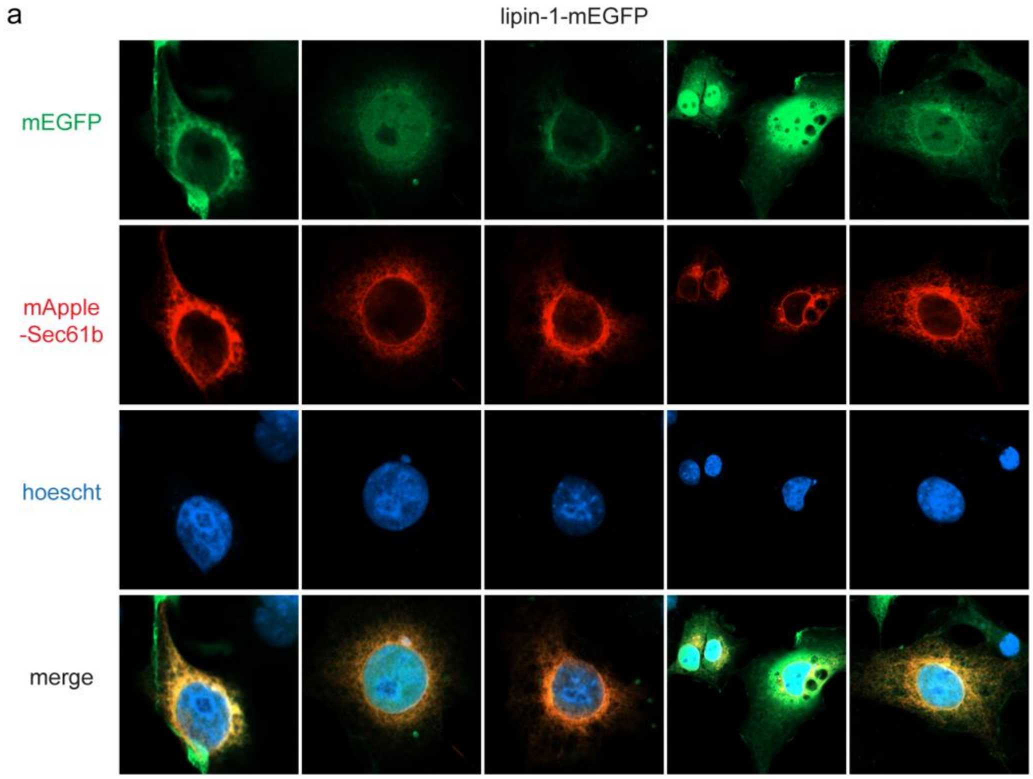
a
lipin-1-mEGFP
mEGFP
mApple
-Sec61b
hoescht
merge

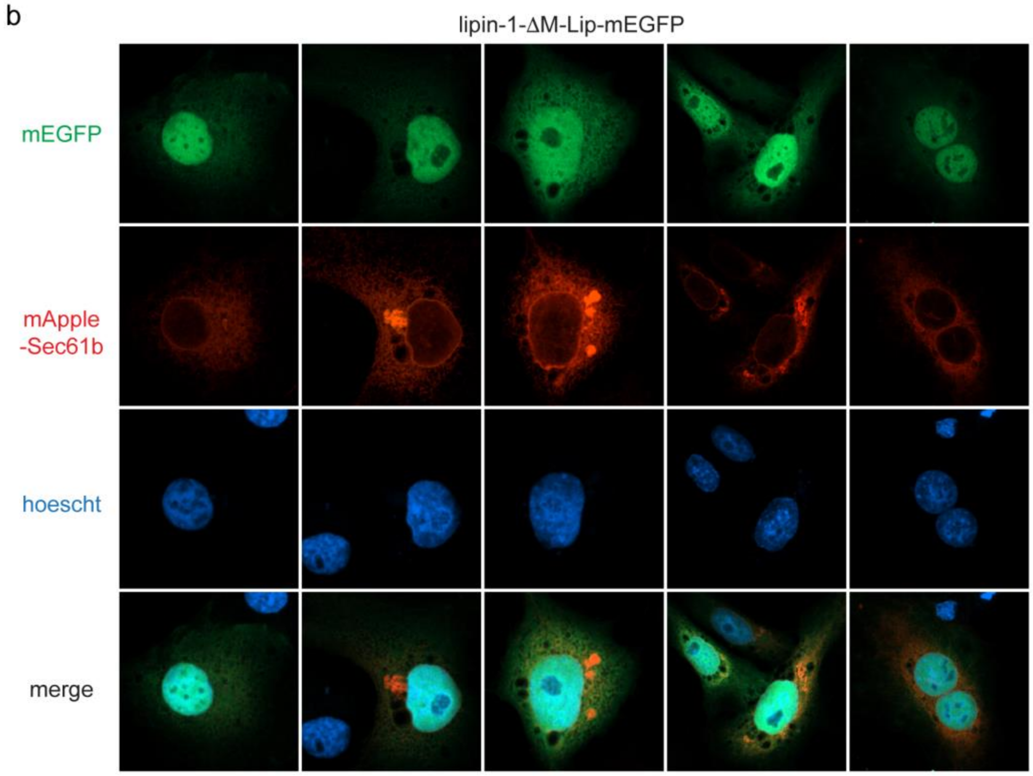
b
lipin-1-ΔM-Lip-mEGFP
mEGFP
mApple
-Sec61b
hoescht
merge

**Supplementary Figure 3.** Confocal microscopy images of Cos-7 cells transiently transfected with monomeric enhanced GFP (mEGFP) fusions of either **(a)** lipin-1 or **(b)** lipin-1 ΔM-Lip (green) and the ER marker mApple-Sec61b (red). Hoechst stain (blue), nucleus.

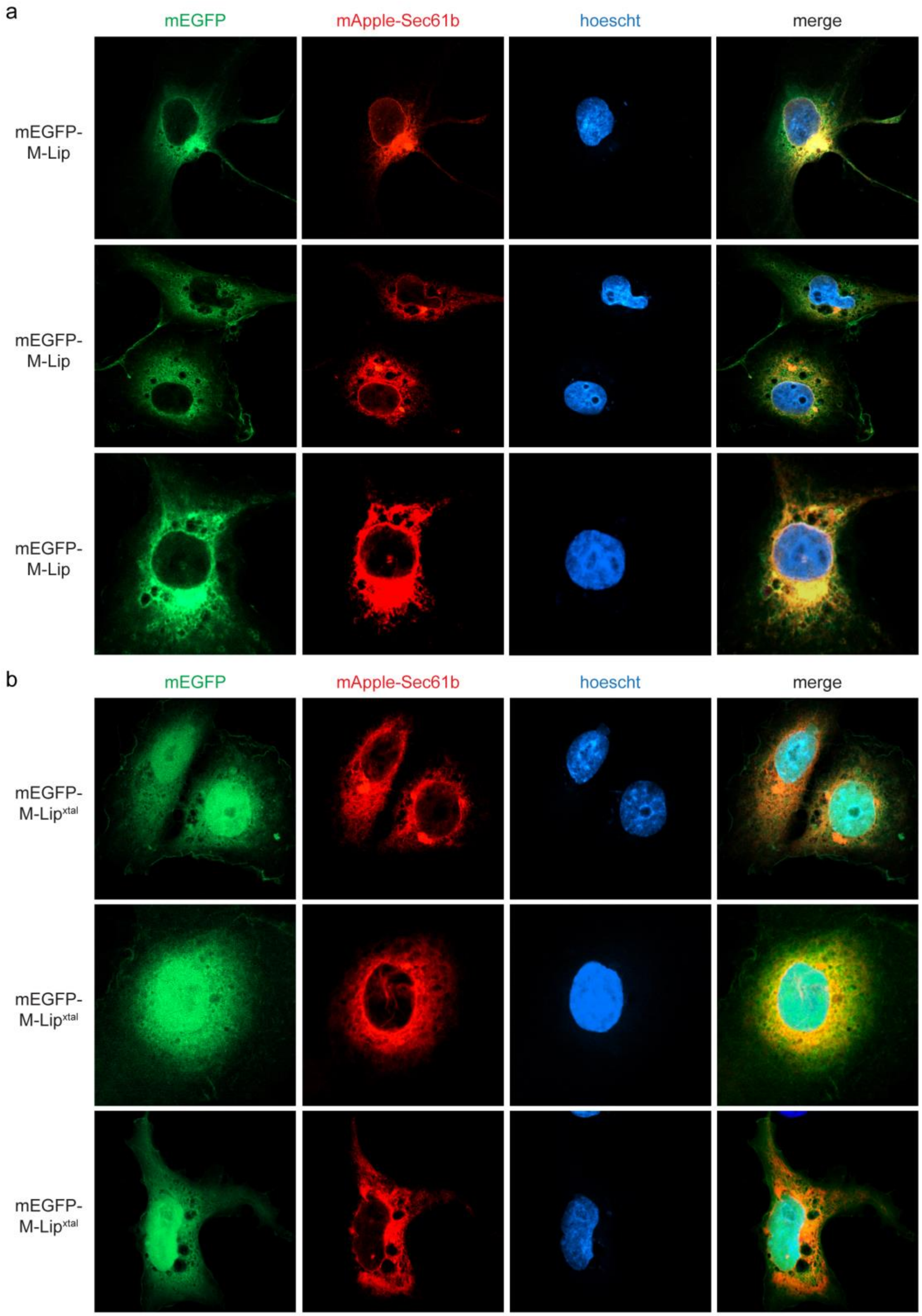


**Supplementary Figure 4.** Confocal microscopy images of Cos-7 cells transiently transfected with monomeric enhanced GFP (mEGFP) fusions of either the **(a)** M-Lip or **(b)** M-Lip$^{xtal}$ domains (green) and the ER marker mApple-Sec61b (red). Hoechst stain (blue), nucleus.

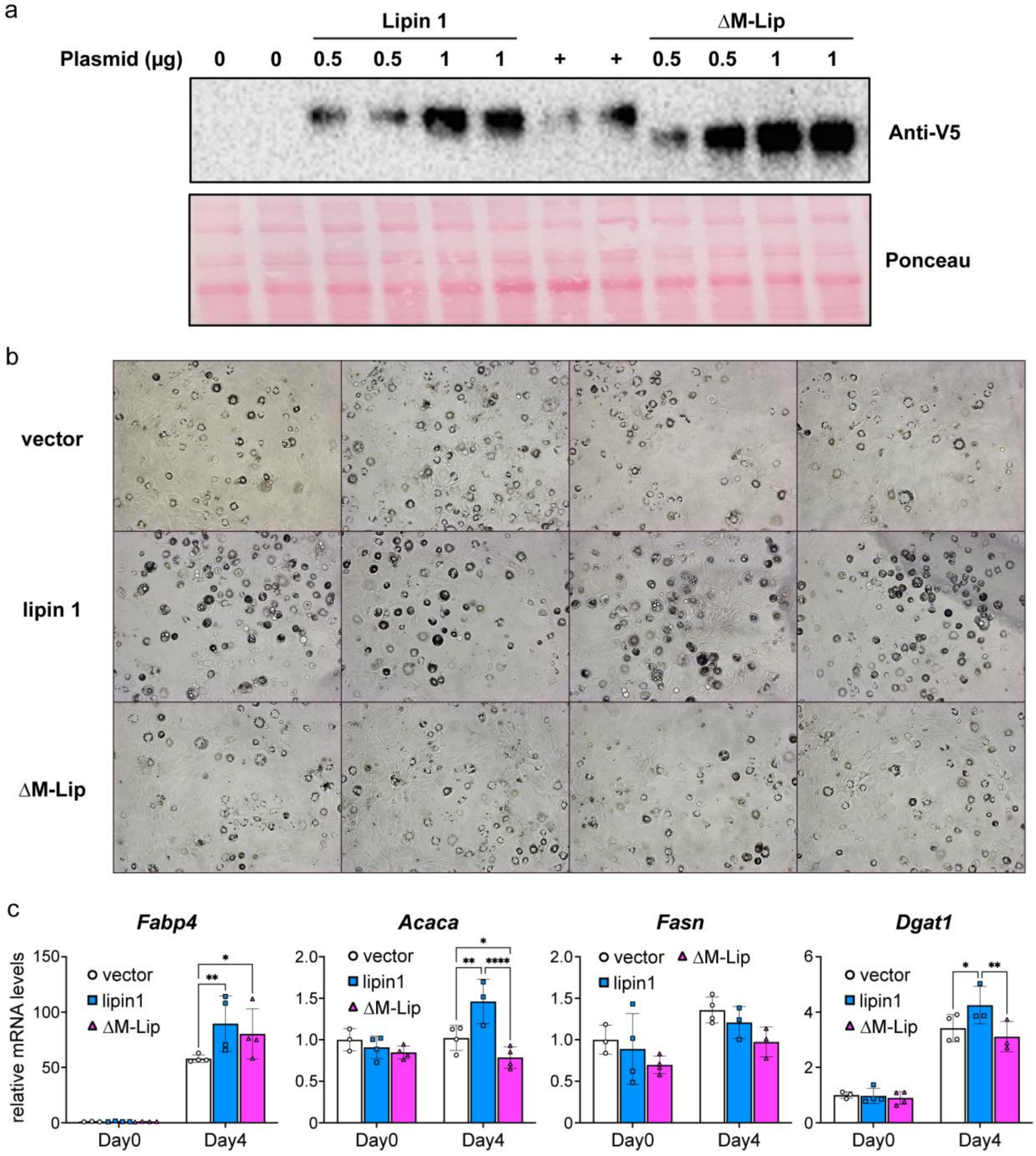


**Supplementary Figure 5. Additional data regarding role of M-Lip in lipin 1 enhancement of adipogenesis.**

**a.** Similar protein levels of wild-type lipin 1 and ΔM-Lip protein expression in 3T3-L1 transfection experiments. For experiments shown, 0.5 μg of each construct was used for transfections. +, recombinant lipin 1 produced in HEK293 cells as control for antibody detection. Total protein loads shown with Ponceau-stained blot.

**b.** Lipid accumulation at intermediate stage (day 5) of 3T3-L1 differentiation in an independent experiment from that shown in Fig. 5b. Bright field illumination shows lipid accumulation as opaque areas within cells (100x).

**c.** Gene expression from an independent experiment to that in Fig. 5c showing expression of genes involved in fatty acid synthesis (*Acaca*, *Fasn*) and triglyceride synthesis (*Dgat1*). *Fabp4*, fatty acid binding protein 4; *Acaca*, acetyl CoA-carboxylase; *Fasn*, fatty acid synthase; *Dgat1*, diacylglycerol acyltransferase 1. Gene expression was analyzed by 2-way ANOVA. *, $p<0.05$; **, $p<0.01$; ***$p<0.001$. n=4 for each group.
